# Cross-linking of the Endolysosomal System Reveals Flotillin Structures and Putative Cargo

**DOI:** 10.1101/2022.01.12.475930

**Authors:** Jasjot Singh, Hadeer Elhabashy, Pathma Muthukottiappan, Markus Stepath, Martin Eisenacher, Oliver Kohlbacher, Volkmar Gieselmann, Dominic Winter

## Abstract

Lysosomes are well-established as the main cellular organelles for the degradation of macromolecules and emerging as regulatory centers of metabolism. They are of crucial importance for cellular homeostasis, which is exemplified by a plethora of disorders related to alterations in lysosomal function. In this context, protein complexes play a decisive role, regulating not only metabolic lysosomal processes, but also lysosome biogenesis, transport, and interaction with other organelles. Using cross-linking mass spectrometry, we analyzed lysosomes and early endosomes. Based on the identification of 5,376 cross-links, we investigated protein-protein interactions and structures of lysosome- and endosome-related proteins. In particular, we present evidence for a tetrameric assembly of the lysosomal hydrolase PPT1 and heterodimeric/- multimeric structures of FLOT1/FLOT2 at lysosomes and early endosomes. For FLOT1-/FLOT2- positive early endosomes, we identified >300 proteins presenting putative cargo, and confirm the latrophilin family of adhesion G protein-coupled receptors as substrates for flotillin-dependent endocytosis.

## INTRODUCTION

Lysosomes, the central lytic organelles of mammalian cells, are of crucial importance for cellular homeostasis. This is underscored by the detrimental consequences resulting from impairment of lysosomal function: mutations in genes encoding lysosomal proteins are causative for a group of around 70 rare and frequently devastating diseases, so-called lysosomal storage disorders (LSDs). Moreover, lysosomal dysfunction has been demonstrated in a number of more common conditions, including neurodegenerative diseases and cancer (Fraldi et al., 2016; Platt, 2018).

In addition to the long-known role of lysosomes in the degradation of intra- and extracellular substrates, more recent findings place them at the center of metabolic signaling. The major player in this context is the mammalian target of rapamycin complex 1 (mTORC1), whose activity is regulated at the lysosomal surface. This regulation is mediated by several protein complexes located in/at the lysosomal membrane, which integrate the activity of major signaling pathways, as well as the concentration of various metabolites (Shin and Zoncu, 2020). Furthermore, protein complexes were shown to play a role in other lysosome-related processes, such as their transport, direct interaction with other cellular compartments, gene regulation, immunity, cell adhesion/migration, and plasma membrane repair (Ballabio, 2019)

For most of these functions, protein-protein interactions at the lysosomal membrane play a decisive role. Nutrient sensing and activation of mTORC1 is regulated by the interaction of at least 30 individual proteins (Liu and Sabatini, 2020), and lysosomal motility is controlled by the reversible association to microtubules through dynein and kinesin by several adaptor/scaffold complexes, such as BLOC1-related complex (BORC) (Cabukusta, 2018). The core feature of lysosomes, their acidic pH, is maintained by the 1.25 MDa vacuolar-type ATPase (V-ATPase) complex, which consists of 35 subunits (17 unique proteins), and catalyzes the transport of protons across the lysosomal membrane (Wang, 2020). Delivery of certain lysosomal proteins is achieved by members of the homotypic fusion and protein sorting (HOPS) (Garg et al., 2011), the class C core vacuole/endosome tethering (Corvet) (Balderhaar, 2013), as well as adaptor protein (AP) complexes, and the endosomal sorting complex required for transport (ESCRT) mediates repair of lysosomal membranes (Skowyra et al., 2018).

Also, for the interactions of lysosomes with other organelles, protein complexes play an essential role. This includes fusion events with cargo delivery vesicles such as endosomes, phagosomes, and autophagosomes (Cheng et al., 2010), exocytosis at the plasma membrane (Reddy et al., 2001), or direct interactions with the endoplasmic reticulum (Levin-Konigsberg et al., 2019), the Golgi apparatus (Hao et al., 2018) peroxisomes (Chu et al., 2015), RNA granules (Liao et al., 2019), and mitochondria (Wong, 2018). The latter facilitates, for example, the exchange of small molecules and was shown to regulate events such as mitochondrial fusion and fission (Ballabio, 2019).

The majority of protein complexes that facilitate these processes are poorly characterized, and novel members/interactors are continuously being identified. Given the central role of lysosomes in metabolic regulation, and the high number of cellular structures they interact with, it is highly likely that several functionally important interactors of lysosomal proteins are still unknown. Although structural data are available for a number of lysosomal luminal proteins and complexes in/at its membrane, three-dimensional information is still lacking for a significant fraction of the lysosomal proteome. The majority of existing structural data originates from crystallography experiments, heavily relying on affinity-purified proteins, or fragments thereof, from pro- or eukaryotic overexpression systems, and crystallization *in vitro*. The applicability of these structures to the *in vivo* situation remains, therefore, in some instances, questionable (Niedzialkowska et al., 2016).

A promising avenue to identify unknown interactions of lysosomal proteins, and to reveal new insights into their structure under physiological conditions, is chemical cross-linking in combination with mass spectrometry-based proteomics (XL-LC-MS/MS) (O’Reilly and Rappsilber, 2018). In cross-linking experiments, a chemical linker forms covalent bonds between certain amino acids such as lysine. In subsequent MS analyses these bonds are identified, providing direct proof for the interaction of proteins within a certain distance constraint, defined by the type of cross-linker (Yu and Huang, 2018). This allows for the identification of protein-protein interactions (PPIs), and hence localization, with high confidence. Compared to other commonly used approaches, such as immunoprecipitation (IP), proximity labeling (Kim and Roux, 2016) or lysosome-enrichment (Muthukottiappan and Winter, 2021), such data provide superior spatial evidence. Furthermore, the distance constraints of the cross-linker can serve as a basis for the molecular modeling of proteins and their complexes. This allows supplementing well-established techniques such as nuclear magnetic resonance (NMR), X-ray crystallography, or cryo-electron microscopy (cryo-EM), compensating for missing/incomplete data, and validating predicted protein structures (Barysz, 2018).

So far, XL-LC-MS/MS experiments have been performed for samples of varying complexity, ranging from individual proteins (Chen, 2010) and multi-subunit complexes (Albanese, 2020; O’Reilly et al., 2020), to whole organelles (Fasci, 2018; Schweppe et al., 2017) and cell/tissue lysates (Chavez et al., 2019; Liu et al., 2015). Dedicated analysis of lysosomal proteins by cross-linking has not been performed to date, which is certainly related to the fact that lysosomal proteins are of low abundance (estimated 0.2 % of cellular protein mass (Itzhak et al., 2017; Valm et al., 2017)).

In the current study, we present the first cross-linking dataset of the endolysosomal compartment of HEK293 cells, applying the MS-cleavable cross-linker disuccinimidyl sulfoxide (DSSO) to lysosome-/early endosome-enriched fractions. We present an interaction map of lysosomal proteins, of which we verify selected PPIs by co-IP and validate/extend existing protein structures. Based on the cross-linking data and computational modeling, we further propose higher-order structures for PPT1 and flotillin assemblies. Finally, by affinity purification and MS analysis of flotillin-positive early endosomes, we investigate the putative cargo of these vesicles.

## RESULTS

### Cross-Linking Mass Spectrometry Analysis of Lysosome Enriched Fractions

In mammalian cells, the majority of lysosomal proteins are of relatively low abundance, and whole-cell XL-LC-MS/MS studies typically cover only a fraction of the lysosomal proteome (Figure S1A). A way to overcome this limitation is lysosome enrichment, which we showed to increase signal intensities for certain lysosomal proteins up to 100-fold relative to whole cell lysates (Singh et al., 2020). Accordingly, we enriched lysosomes by superparamagnetic iron oxide nanoparticles (SPIONs, Figure 1A), and established cross-linking conditions for lysosome-enriched fractions utilizing the MS-cleavable cross-linker DSSO (Kao et al., 2011). Due to a limited membrane permeability of DSSO (Singh et al., 2021), we cross-linked lysosomes both in an intact (IT) and disrupted (DR) state, and determined optimal reaction conditions by silver staining and western blot (Figure S1B, C). Subsequently, we enriched lysosomes from 384 plates of HEK293 cells across three biological replicates, and assessed lysosomal intactness, recovery, and enrichment (Figure 1B, 1C). Using a non-cross-linked fraction of each sample, we acquired an LC-MS/MS reference dataset. In total, we identified 4,181 proteins, of which 474 were assigned the term “lysosome” based on GO terms and UniProt classifiers in >3 runs, indicating an excellent performance of lysosome enrichment (Akter, 2020). To assess the quantitative distribution of lysosomal proteins in our samples, we utilized these data to estimate absolute protein abundances by intensity-based absolute quantification (iBAQ) (Schwanhausser et al., 2011). This revealed a three-fold overrepresentation of lysosomal protein abundance relative to the whole dataset (Figure 1D).

**Figure 1:**
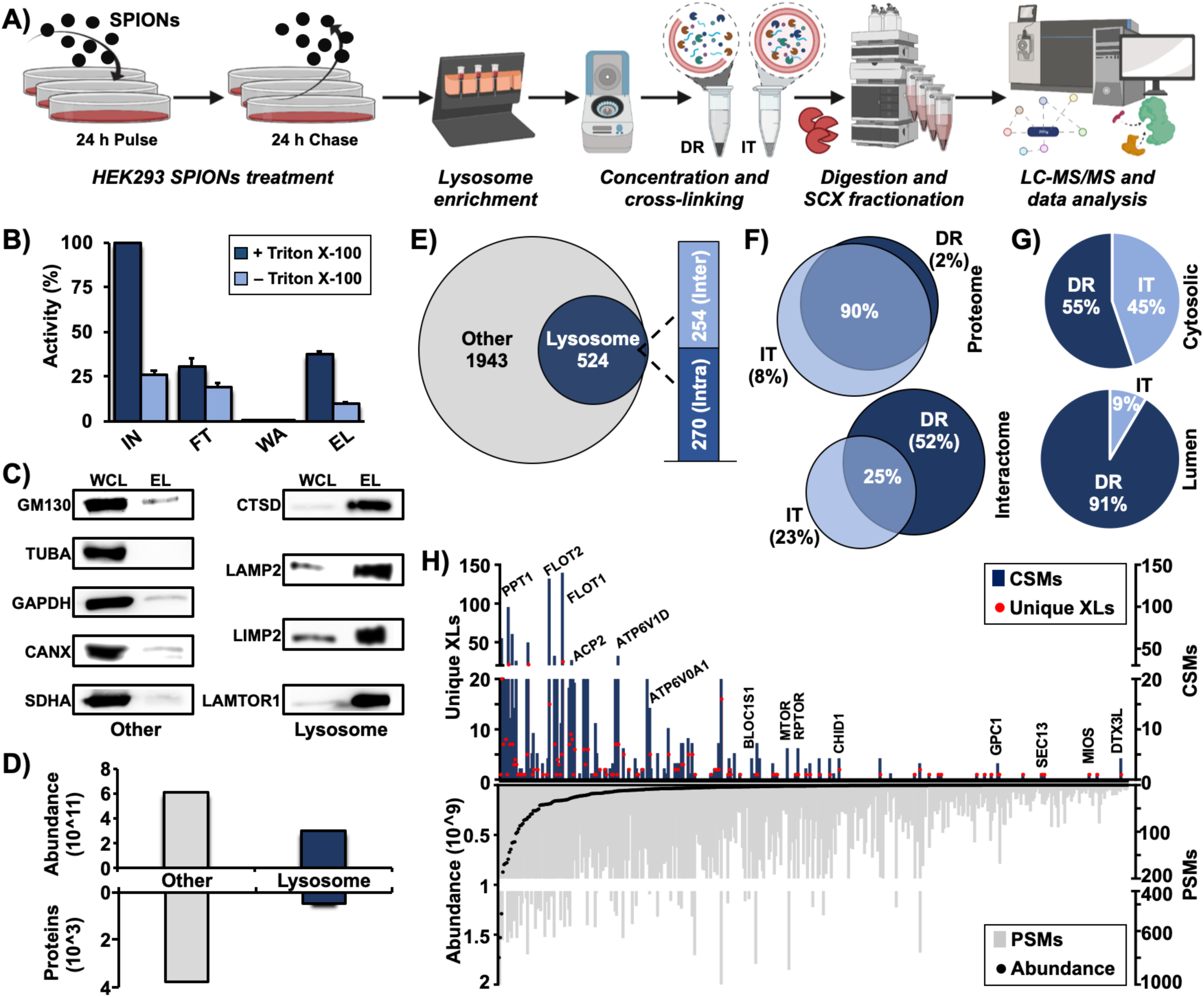
Cross-linking mass spectrometry analysis of lysosome-enriched fractions. **A)** Experimental workflow for the XL-LC-MS/MS analysis of lysosome-enriched fractions. **B)** Normalized β-hexosaminidase activities for individual fractions from lysosome enrichment by SPIONs. Shown are average values (n=3, +STDEV). **C)** Western blot analysis of lysosome-enriched fractions for contamination by other organelles. Lysosome: lysosomal proteins (CTSD, LAMP2, LIMP2, and LAMTOR1). Other: Golgi-apparatus (GM130), cytoskeleton (TUBA), cytosol (GAPDH), endoplasmic reticulum (CANX), mitochondria (SDHA). **D)** Summed iBAQ abundances for proteins identified in lysosome-enriched fractions in ≥ 3 replicates. **E)** Classification of unique cross-linked residue pairs. **F)** Proteins detected in non-cross-linked lysosome enriched fractions (proteome) and unique lysosomal cross-linked residue pairs (interactome) for DR and IT samples. **G)** Localization of CSMs for 68 lysosomal proteins cross-linked in the DR and IT state. Cytosolic: proteins located at the cytosolic face of lysosomal membrane; Lumen: lysosomal luminal proteins. **H)** Correlation of cross-link identification and protein abundance for lysosomal proteins. CSMs and PSMs represent summed values of the analysis of 6 replicates (DR and IT). **Abbreviations:** SPIONs: superparamagnetic iron oxide nanoparticles; DR: disrupted; IT: intact; SCX: strong cation-exchange; IN: input; FT: flow through; W: wash; EL: eluate; WCL: whole cell lysate; iBAQ: intensity based absolute quantification; XL: cross-links; CSMs: cross-link spectral matches; PSMs: peptide spectral matches. **See also**: Figure S1.

We cross-linked lysosome enriched fractions in both the IT and DR state, followed by their proteolytic digestion, strong cation-exchange (SCX) peptide fractionation, and analysis by LC-MS/MS (Figure 1A). Analysis of the XL-LC-MS/MS dataset with XlinkX (Klykov et al., 2018) resulted in the assignment of 6,580 cross-link spectral matches, originating from 4,294 cross-linked peptides. Out of the 2,467 unique residue-to-residue cross-links, 524 identifications (270 intra links between different residues of the same protein and 254 inter-links between two different proteins) originated from 111 proteins assigned to the lysosomal compartment (Figure 1E, Figure S1D-H). Interestingly, only 25 % of cross-links were found both in the IT and the DR state, while the latter contributed a larger fraction to the dataset, further demonstrating the limited membrane permeability of DSSO (Figure 1F). A similar distribution was observed for the cross-links identified for non-lysosomal proteins contained in the dataset (Figure S1D). Strikingly, while cross-link spectral matches (CSMs) of cytosolic proteins were almost equally distributed between both conditions, 91 % of the dataset’s CSMs assigned to lysosomal luminal proteins were identified in samples cross-linked in the DR state (Figure 1G).

As lysosomal proteins are expressed at a dynamic abundance range encompassing three orders of magnitude (Akter, 2020), we further correlated CSMs, peptide spectral matches (PSMs), and iBAQ values. Even though higher abundant proteins tended to yield more CSMs, we did not identify a strong correlation between cross-link identification and protein abundance, showing that our dataset also covered proteins of low expression levels (Figure 1H, Figure S1I). When we compared the average iBAQ abundance for proteins involved in intra- and inter-links, we observed a tendency towards the identification of more intra-links in higher abundant proteins, which was less pronounced for lysosomal proteins (Figure S1J, K).

Finally, we investigated the distribution of CSMs across a short-list of lysosomal proteins/complexes (Muthukottiappan and Winter, 2021). Most categories showed an equal distribution between DR and IT samples, with the exception of proteins involved in lysosomal substrate degradation (93 %), heat shock proteins (70 %), and annexins (75 %), for which more cross-links were annotated in the DR sample (Figure S1L).

### Characterization of the Human Lysosomal Interactome

Based on all inter-links contained in the dataset, we constructed a network of 1,008 proteins engaged in 1,023 interactions, of which 254 involved lysosomal proteins (Figure 2A, Figure S2A-C). Comparison to known interactions revealed an overlap of ∼30 %, confirming the validity of our dataset. While 34 % of interactions of non-lysosomal proteins were included in STRING, only 26 % of potential lysosomal PPIs have been reported previously (Figure 2B). We classified lysosomal PPIs based on the interacting subcellular compartment, revealing an overrepresentation of nuclear and cytoplasmic/cytoskeletal proteins (Figure 2C). With respect to lysosomal and lysosome-associated proteins, we identified the highest numbers of PPIs for the V-ATPase, the flotillins, mTORC1, and the syntaxins (Figure 2D, E, Figure S2D, E).

**Figure 2:**
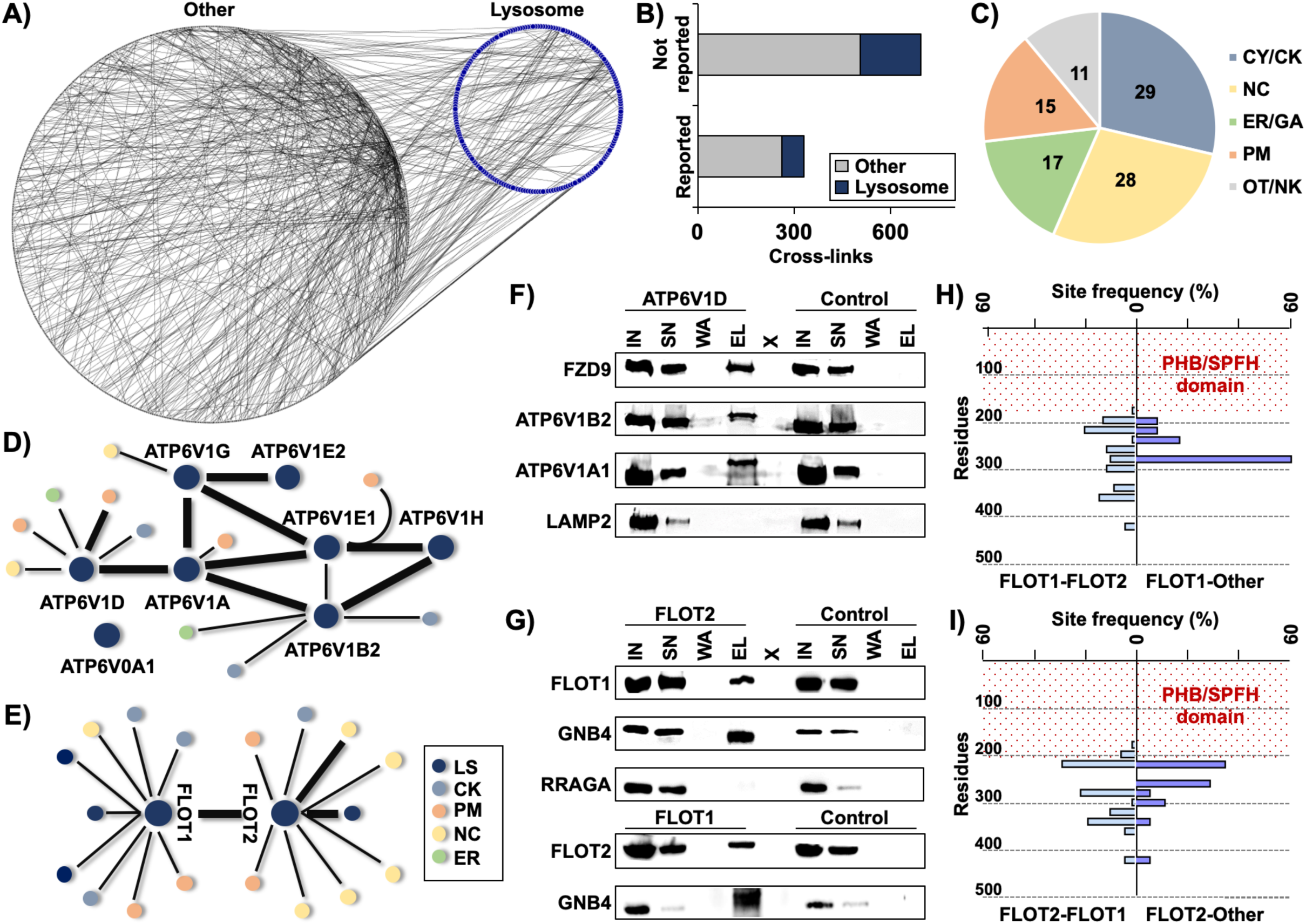
The human lysosomal interactome. **A)** PPIs identified in the XL-LC-MS/MS dataset. Lysosomal proteins (blue dots), non-lysosomal proteins (grey dots), and PPIs (grey lines) are indicated. **B)** Matching of PPIs to the STRING database. **C)** Numbers of proteins from distinct subcellular localizations interacting with lysosomal proteins. **D/E)** Interaction networks of the V-ATPase (D) and the flotillin (E) complex. **F)** Co-IP of ATP6V1D and FZD9. ATP6V1B2 and ATP6V1A1 are known members of the V-ATPase complex; LAMP2 is a lysosomal membrane protein. Control: empty beads. **G)** Co-IPs of FLOT1, FLOT2, and GNB4. RRAGA is a lysosomal membrane-associated protein. Control: empty beads. **H/I)** Site frequency distribution for identified FLOT1 (H) and FLOT2 (I) cross-links. Site frequency represents the percentage of cross-links detected in bins of 20 residues each. The region indicated by red dots represents the PHB domain. **Abbreviations:** CY: cytoplasm; CK: cytoskeleton; NC: nucleus; ER: endoplasmic reticulum; GA: Golgi apparatus; PM: plasma membrane; LS: lysosome; OT: others; NK: not known; IN: input; SN: supernatant; W: wash; EL: eluate; X: empty lane. **See also:** Figure S2.

For the V-ATPase, we detected most PPIs for the D subunit of its soluble V1 part (ATP6V1D). This may be related to the capability of V1 to dissociate from the lysosomal membrane-embedded V0 part, which was shown to be involved in the regulation of V-ATPase activity (Maxson and Grinstein, 2014), exposing the D subunit to interactions. The cross-link of ATP6V1D with frizzled 9 (FZD9), a member of the WNT signaling pathway, sparked our interest. FZD9 was shown to be sorted to late endosomes/lysosomes after its internalization by endocytosis (Grainger et al., 2019), and the ATP6AP2 subunit of the V-ATPase was reported to interact with FZD8 (Cruciat et al., 2010). We therefore investigated the validity of this PPI by co-IP, confirming both the observed interaction of ATP6V1D with FZD9, as well as interaction with other subunits of the complex (Figure 2F).

We further investigated the interaction of FLOT1, FLOT2, and GNB4, for which we identified an inter-link with FLOT2. We were able to co-IP FLOT1 and FLOT2, which are known to form heterooligomers (Babuke et al., 2009), as well as GNB4 with its direct interactor FLOT2 as well as with FLOT1, indicating binding of GNB4 to FLOT1/FLOT2 heteromeric assemblies (Figure 2G).

As we observed 18 different PPIs for FLOT1 and FLOT2, we further investigated their distribution across both proteins. While FLOT1/FLOT2 inter-links were detected across most of the regions predicted to form a helical structure (amino acids 193-365 and 213-362 for FLOT1 and FLOT2, respectively) (Rivera-Milla et al., 2006), the interaction with other proteins occurred almost exclusively in confined sections of < 100 amino acids (Figure 2H, I). While the equal distribution of inter-links shows that no sequence-dependent bias towards cross-link detectability exists, localization of the majority of PPIs to a distinct part of the proteins suggests the presence of FLOT1/FLOT2 interaction hotspots.

### Structural Integration of Cross-Linker Distance Constraints Suggests a Tetrameric Assembly of PPT1 *In Vivo*

Cross-links between different amino acids provide distance information (the length of DSSO links is ∼35 Å) that is helpful to validate (or infer) protein structures (Kastritis et al., 2017). Initially, we used TopoLink (Ferrari et al., 2019) to match 161 unique cross-links to the resolved structures of 34 lysosomal and lysosome-associated proteins, confirming the validity of our dataset. The remaining 64 cross-links assigned to lysosomal proteins could not be integrated, as the respective regions have not been resolved yet. We matched these cross-links either to homology models based on available PDB structures from other organisms using SWISS-MODEL (Waterhouse et al., 2018), or to predicted AlphaFold models which became available during the preparation of this manuscript (Jumper et al., 2021) (Figure 3A, Figure S3).

**Figure 3:**
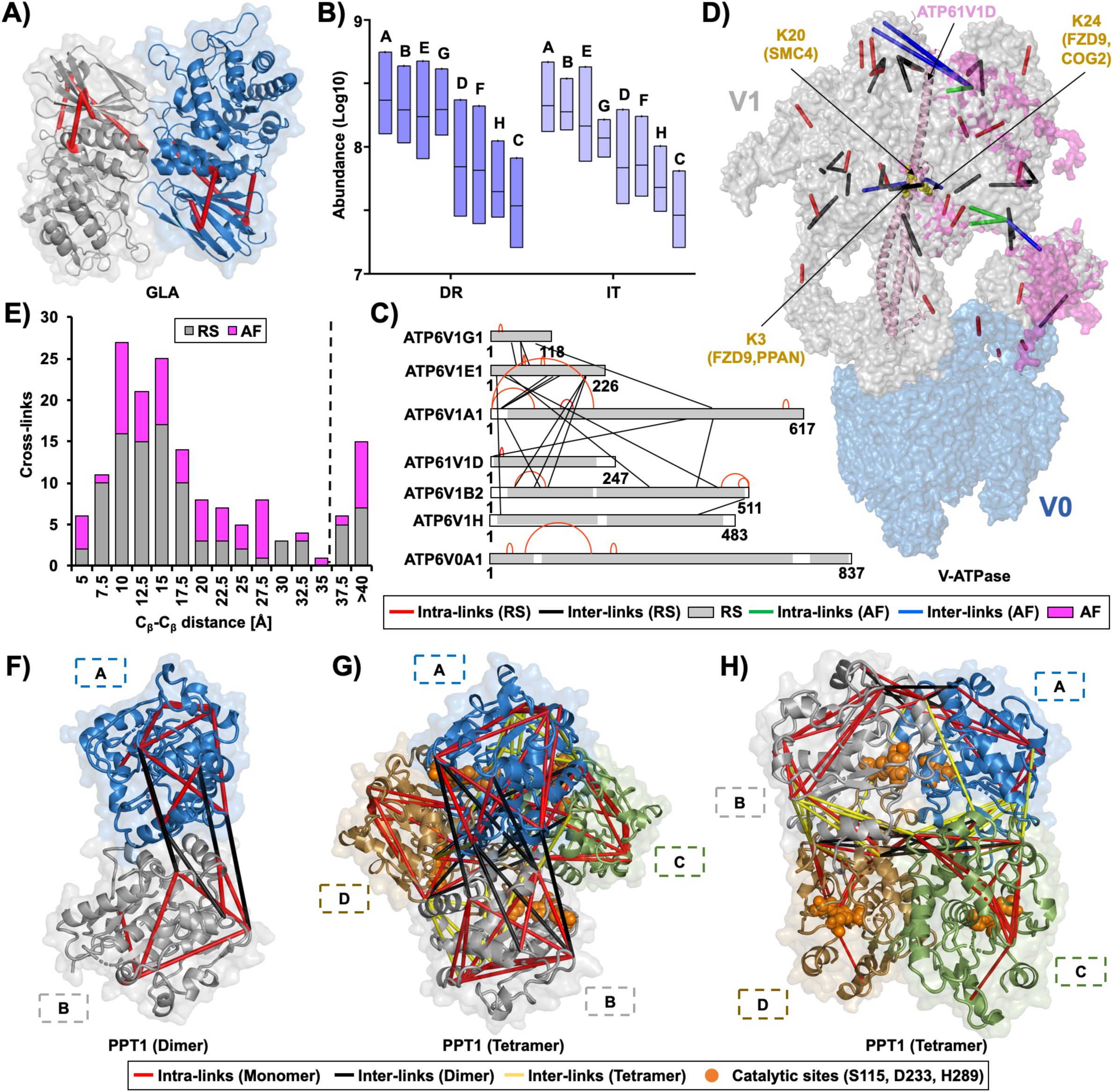
Cross-link-based structural refinement of the V-ATPase and PPT1 complex. **A)** Matching of cross-links to the crystal structure of lysosomal alpha-galactosidase A (GLA). **B)** Average iBAQ abundances for individual V-ATPase subunits based on the analysis of lysosome-enriched fractions in a non-cross-linked state. **C)** Cross-links identified for individual V-ATPase subunits. Structurally resolved regions are colored grey, unresolved regions white. **D)** Refined structure of the V-ATPase complex. Identified cross-links were integrated in the V-ATPase cryo-EM structure (Wang, 2020). Missing regions were supplemented with predicted structures from AlphaFold based on the identified cross-links. **E)** Distances for mapped cross-links to crystal structures and AlphaFold models for lysosomal proteins. Bin size: 2.5 Å. **F)** Reported homodimeric human PPT1 structure (PDB: 3GRO). **G/H)** Tetrameric PPT1 model representing the most favorable energetic state and fulfilling DSSO’s distance constraints for all 18 cross-links. A, B, C, and D indicate individual subunits. **Abbreviations:** DR: disrupted; IT: intact; RS: resolved structure; AF: AlphaFold. **See also:** Figure S3.

The complex for which we identified the highest number of cross-links was the V-ATPase, a 35-mer assembly of 17 unique proteins (Wang, 2020), of which 13 were covered in our dataset. While the soluble V1 part (eight different proteins) yielded 26 cross-links, only three were identified for the membrane-embedded V0 section (seven different proteins). After confirmation of the V1 part’s correct stoichiometry through the individual subunit’s iBAQ values in the non-cross linked dataset (A3B3E3G3D1F1C1H1) (Figure 3B), we mapped the identified cross-links to a recently published structure determined by cryo-EM (Wang, 2020). Out of the 29 unique cross-links identified, 21 could readily be integrated into the published structure, while 8 originated from regions that were so-far structurally not resolved (Figure 3C). For these sections, we integrated the predicted full-length protein models by AlphaFold, aligning them with the published V-ATPase sub-unit structures in the complex, based on the identified cross-links (Figure 3D). In this combined model, we cover ∼95 % of the V-ATPase sequence, and 90 % of cross-links fulfil DSSO’s distance constraints. The remaining three cross-links were inter-links of the A and E subunit of the V1 part, which undergo conformational changes during ATP hydrolysis (Wang, 2020).

With respect to mTORC1, we identified most cross-links for proteins related to Ragulator, a lysosomal membrane-associated complex that is crucial for mTORC1 activity (Shin and Zoncu, 2020). For the N-terminal region of LAMTOR1, we identified three cross-links of the same lysine residue with different amino acids of LAMTOR3. Two of them violated DSSO’s distance constraints relative to the crystal structure obtained in the presence of the Ragulator-associated RAG GTPases (De Araujo, 2017), indicating cross-linking of an alternative state, possibly representing Ragulator in the absence of RAG GTPases (Figure S3J). We further identified three intra-links from RRAGA that originated from the same lysine residue at its C-terminus, cross-linked to three different amino acids (Figure S3K. This could be related to a high flexibility of this region of the protein, which is in accordance with the fact that it could not be covered in a previous crystallization study (Su et al., 2017).

Of the 161 lysosomal cross-links identified, 21 exceeded DSSO’s distance constraint (Figure 3E). Surprisingly, nine of them originated from PPT1, a member of the palmitoyl protein thioesterase family. The current PPT1 crystal structure (3GRO) resembles a homodimeric assembly of two identical subunits (Figure 3F). Strikingly, while all 11 PPT1 intra-links fulfilled the distance constraints of this structure, the inter-links exceeded them, indicating the possibility of an alternative oligomerization state. This possibility is in line with a previous study, detecting the maximal enzymatic activity of PPT1 at a complex size of >100 kDa (Lyly et al., 2007). We therefore used HADDOCK to perform restraint-based docking for the monomeric subunits of PPT1, extracted from 3GRO. This resulted in the prediction of a tetrameric PPT1 model, which fulfills the distance constraints for all 18 PPT1 cross-links (Figure 3G, H).

### Proposal of a Heterodimeric FLOT1/FLOT2 Model Featuring Extended Alpha-Helical Domains

We identified the highest number of cross-links for the two members of the flotillin family, FLOT1 and FLOT2, which were also overrepresented in the proteomic dataset of the lysosome-enriched fraction (Figure 1H). FLOT1 and FLOT2 are lipid raft-associated proteins, which are present in nearly every type of vertebrate cell, and are highly conserved among organisms (Rivera-Milla et al., 2006). We confirmed their co-enrichment with lysosomes by western blotting (Figure 4A) and co-localization by immunostaining (Figure 5A), which is in agreement with previous EM studies detecting FLOT1 at the lysosomal surface (Kaushik et al., 2006; Kokubo et al., 2003). It is well-established, that FLOT1 and FLOT2 form heterodimers with a 1:1 stoichiometry (Frick et al., 2007), but only partial structural information is available from NMR analyses of the N-terminal region of mouse Flot2, as purification of the full-length proteins is problematic (Dempwolff et al., 2016).

**Figure 4:**
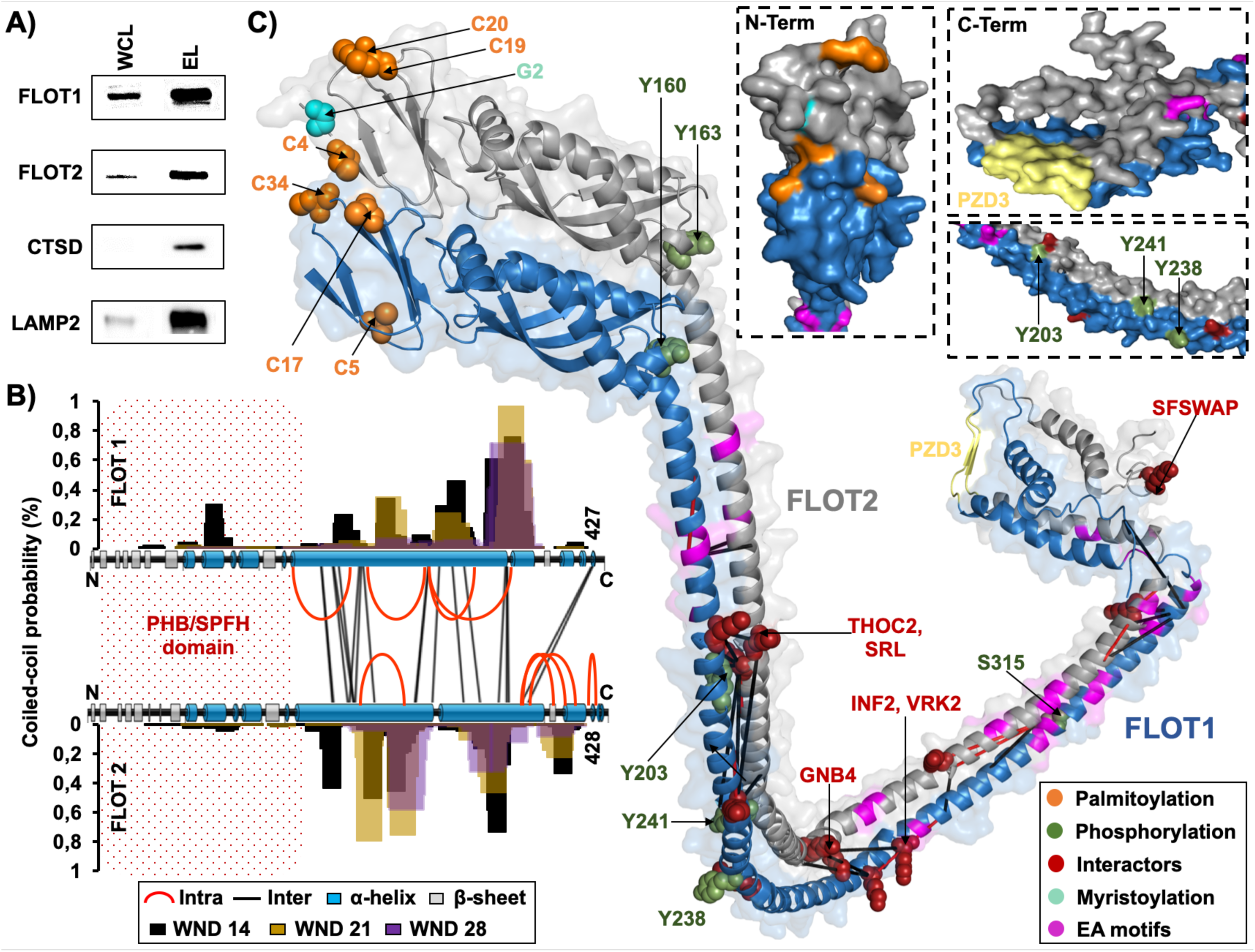
Proposed model for a heterodimeric FLOT1-FLOT2 complex. **A)** Western blot analysis of lysosome-enriched fractions. CTSD is a lysosomal luminal and LAMP2, a lysosomal transmembrane protein. **B)** Identified cross-links, predicted secondary structures (PSIPRED), and coiled-coil probabilities (PCOILS) for FLOT1 and FLOT2. **C)** Heterodimeric FLOT1/FLOT2 interaction model representing the lowest energy state and satisfying the distance constraints of all cross-links. The model was generated using HADDOCK based on predicted monomeric AlphaFold structures. **Abbreviations:** WCL: whole cell lysate; EL: eluate; PHB: prohibitin homology domain; SPFH: stomatin/PHB/flotillin/HflK/C domain; WND: window; EA: glutamic acid/alanine; PZD3: postsynaptic density protein-95/discs large/zonula occludens-1. **See also:** Figure S4.

**Figure 5:**
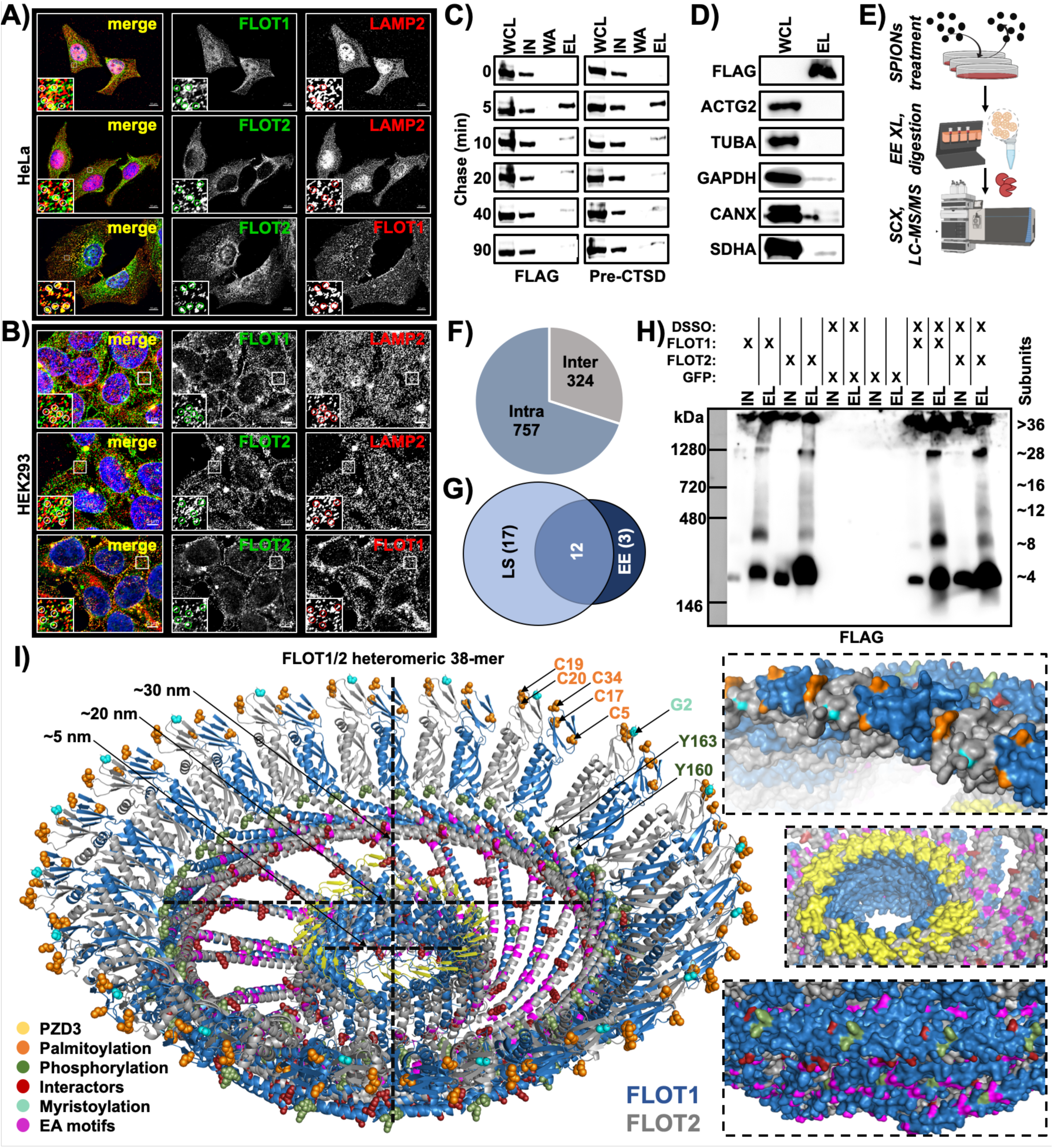
Investigation of higher-order flotillin assemblies in lysosome- and early endosome-enriched fractions. **A/B)** Immunostaining of HeLa and HEK293 cells for FLOT1, FLOT2, and the lysosomal marker LAMP2. **C)** Analysis of FLOT1/FLOT2 in early endosomes. SPIONs pulse treatment time course followed by endosome enrichment, chase times are indicated. Pre-CTSD serves as a marker for early endosomes. **D)** Western blot analysis of early endosome fractions with marker proteins for different subcellular compartments: Cytosol (ACTG2 and GAPDH), cytoskeleton (TUBA), endoplasmic reticulum (CANX), and mitochondria (SDHA). **E)** Experimental workflow for early endosome enrichment and XL-LC-MS/MS analysis. **F)** Overview of early endosome cross-linking dataset. **G)** Overlap of unique FLOT1/FLOT2 cross-links for XL-LC-MS/MS analyses of early endosome- and lysosome-enriched fractions. **H)** Western blot analysis of BN-PAGE separated FLAG-IP eluates with/without cross-linking by DSSO. IN/EL refers to fractions of FLAG-IP. **I)** Nominated lowest energy model for the FLOT1/FLOT2 heteromeric 38-mer. **Abbreviations:** DSSO: disuccinimidyl sulfoxide; M: marker; IN: input; WA: wash; EL: eluate; WCL: whole cell lysate; SPIONs: superparamagnetic iron oxide nanoparticles; SCX: strong cation-exchange; LS: lysosome; EE early endosome; EA: glutamic acid/alanine; PZD3: postsynaptic density protein-95/discs large/zonula occludens-1. **See also:** Figure S5.

Our dataset contains 29 unique cross-links for FLOT1 and FLOT2, including 11 intra- and 22 inter-links. In a first step, we predicted individual secondary structures for both FLOT1 and FLOT2 (Figure 4B) using PSIPRED (Jones, 1999) For the N-termini, this resulted in a cluster of beta-sheets, which is in accordance with their sequence homology to the stomatin/PHB/flotillin/HflK/C (SPFH) domains and the Flot2 NMR structure. The middle section features an extended α-helical region that is interrupted once in the case of FLOT2, while the C-terminus forms one beta-sheet and several short helices for both proteins. We further calculated coiled-coil probabilities for the helical regions with PCOILS (Zimmermann et al., 2018) with window sizes of 14, 21, and 28 amino acids (Figure 4B). Dependent on the region of both proteins, windows of 14 and 28 amino acids delivered the best results, with slightly different patterns for FLOT1 and FLOT2. Matching of our data to full-length structural models from AlphaFold showed an excellent agreement and all identified cross-links confirmed the predicted structures.

Subsequently, we built heterodimeric models using ColabFold (Mirdita et al., 2021), containing closely aligned highly similar structures for both flotillins. In particular, they feature a globular N-terminal region consisting of SPFH domains (residues 1-162 of FLOT1 and FLOT2), a central linear α-helical region (residues 163-341 of FLOT1 and 163-350 of FLOT2), and a C-terminal α-helical coiled-coil structure (residues 342-427 of FLOT1 and 351-426 of FLOT2). The SPFH domains of both FLOT1 and FLOT2 present with antiparallel β-sheets, with six repeats each, and four partially exposed α-helices, forming an ellipsoidal-like globular domain (Figure 4C, Figure S4). Based on the fact that the major helices were not interrupted, and that the C-terminus was in its most compact state, we selected model four, which was also supported by all FLOT1/FLOT2 cross-links detected (Figure 4C, Figure S4C).

We integrated several structural features which are known for both flotillins into our model (Figure 4C). The *S*-palmitoylation and *N*-myristoylation sites, which are crucial for membrane-association of both flotillins (Neumann-Giesen et al., 2004), are located at the SPFH domain’s membrane interfaces. The tyrosine phosphorylation sites, which were shown to be crucial for flotillin-mediated endocytosis and FLOT1/FLOT2 interaction (Bach and Bramkamp, 2015; Riento et al., 2009), are surface-exposed and in proximity to a basic motif (HQR) on the respective other flotillin. Moreover, the PDZ3 domains of both flotillins strongly co-localize, forming a combined feature, and the EA-rich motifs, which were predicted to mediate flotillin oligomerization (Rivera-Milla et al., 2006; Solis et al., 2007), are distributed along the length of the central α-helical region (Kwiatkowska, 2020). Interestingly, the putative interaction hotspot (Figure 2 H, I) locates around the major bend observed in this structure, containing one tyrosine residue (Y238/Y241) in its center. These residues are located in highly conserved sequence motifs (A-X-A-X-L-A-pY-X-L-Q with X: [D/Q or E/Q]), possibly presenting a regulatory switch for FLOT1/FLOT2 PPIs.

### Flotillins Assemble in Similar Higher Order Structures at Lysosomes and Endosomes

While FLOT1 was detected previously at the cytosolic face of lysosomes by EM (Kaushik et al., 2006; Kokubo et al., 2003) assemblies of both FLOT1 and FLOT2 were only shown for the plasma membrane or early endosomes, where they play a role in clathrin-independent endocytosis (Gorbea et al., 2010; Stuermer et al., 2001). In line with these findings, we detected FLOT1 and FLOT2 to partially co-localize with the plasma membrane and lysosomes of HeLa and HEK293 cells (Figure 5A, B). In these analyses, we also observed numerous FLOT1/FLOT2 punctae, which did not co-localize with lysosomes, presenting putative FLOT1/FLOT2-positive endosomes.

In order to elucidate if the structural assembly of FLOT1 and FLOT2 differs between early endosomes and lysosomes, we performed cross-linking of early endosome-enriched fractions. To increase the number of FLOT1-/FLOT2-positive endosomes, we co-transfected cells with FLAG-tagged FLOT1 and FLOT2, as their overexpression was shown to increase the number of FLOT-positive endosomes (Babuke et al., 2009; Frick et al., 2007). After confirmation of their correct localization (Figure S5A, B), we established SPIONs pulse-chase conditions for the enrichment of early endosomes (Figure 5C). Analysis of the early endosome-enriched fraction by western blot and LC-MS/MS verified the presence of marker proteins such as EEA1, the clathrin chains CLTA, CLTB, and CLTC, as well as the RAB-GTPases RAB5, RAB11, and RAB14. Furthermore, we detected FLOT1 and FLOT2, while markers for other organelles were depleted (Figure 5D). We then performed enrichment of early endosomes from FLOT1/-FLOT2-FLAG overexpressing HEK293 cells, established their cross-linking followed by SCX fractionation and investigated them by XL-LC-MS/MS (Figure 5E, Figure S5C-F). Importantly, western blot analysis of FLOT1/FLOT2 aggregation in response to different amounts of DSSO showed that the concentration we used for cross-linking of lysosome-enriched fractions does not result in over-cross-linking (Figure S5D), further supporting the validity of these data. In total, we identified 1,081 cross-links from 414 unique proteins (Figure 5F, Figure S5G). This early endosome dataset contains 15 unique cross-links for FLOT1 and FLOT2, which all matched to our predicted FLOT1-FLOT2 heterodimer within DSSO’s distance constraints, and from which 80 % overlapped with the cross-linking dataset from lysosome-enriched fractions (Figure 5G). These data indicate that early endosome- and lysosome-localized flotillins assemble in a similar way.

It has been shown previously, that flotillins form higher-order assemblies (Solis et al., 2007; Stuermer et al., 2001). We therefore performed blue native (BN)-PAGE experiments to investigate the size distribution of FLOT1/FLOT2 structures in a native and a cross-linked state (Figure 5H, Figure S5H). These analyses revealed that a significant amount of both FLOT1 and FLOT2 migrates at a range corresponding to a tetrameric assembly, while smaller fractions migrated at sizes consistent with higher-order structures exceeding 1 MDa. The cross-linking samples, which were generated with the same reaction conditions as the early endosome and lysosome experiments, presented with the same complex sizes, indicating that the cross-link data represent the native state.

Based on the combined 32 unique cross-links from the lysosome- and endosome-cross-linking datasets, we further investigated possible structures for higher-order assemblies. All of our attempts to model the tetrameric structure did not lead to a plausible outcome. We further addressed the structures exceeding 1 MDa, which were stabilized by cross-linking with DSSO (Figure 5H). We utilized the structure of rat major vault protein (Mvp), which assembles into the 3.7 MDa rat liver vault (PDB: 4V60), as template for a higher-order hetero-oligomeric FLOT1/FLOT2 model, as Mvp shares several structural properties with flotillins. A key feature in this context are the N-terminal PHB and the 42-turn-long cap-helix domains, which are crucial for stabilizing the particle (Tanaka, 2009). We used the rat liver vault structure as a template for building a model based on FLOT1/FLOT2 hetero-oligomers, proposing a 38-mer structure, which could represent one of many possible higher-order assemblies of the FLOT1/FLOT2 heterodimer (Figure 5I). Important features of this structure are the exposed ring of palmitoylation/myristoylation sites, the central arrangement of PDZ domains, and the two rings of phosphotyrosine (pY) residues. While the pY sites known for FLOT1/FLOT2 interaction (Y160/Y163) are located at the inside of the structure, the sites possibly involved in PPI regulation (Y238/Y241) are located at its outside, making them accessible to kinases even after formation of the higher-order structure.

### Analysis of Flotillin-Endosome Cargo Reveals an Overrepresentation of Membrane Proteins and Receptors

Flotillins have been proposed as defining structural components of an endocytic pathway independent of clathrin and caveolin (Frick et al., 2007; Glebov et al., 2006) and were shown to co-localize with early endosomes (Gorbea et al., 2010). In agreement with these findings, we detected their co-enrichment with endosomes (Figure 5C, D). To investigate the putative cargo of FLOT1-/FLOT2-positive early endosomes, we combined early endosome enrichment from HEK293 cells overexpressing FLAG-tagged FLOT1/FLOT2 and immunoprecipitation of FLAG-positive intact vesicles (Figure 6A). Subsequently, we analyzed the resulting fractions by label-free quantification (Figure S6A-D). Based on 5,089 protein groups quantified across all conditions, we were able to define protein populations that were enriched/depleted in FLOT1-/FLOT2-positive endosomes relative to the total cellular pool of early endosomes (Figure 6B). Importantly, the FLOT1-/FLOT2-depleted population of early endosomes contained the clathrin chains CLTA and CLTC, EEA1, and the endosome-related GTPases RAB5 and RAB11, confirming a separation of clathrin- and flotillin-containing early endosomes. We confirmed these findings by co-immunostaining of EEA1 with endogenous and overexpressed FLOT1/FLOT2 in HeLa and HEK293 cells, showing that FLOT1-/FLOT2-positive vesicles present a distinct population from EEA1-positive endosomes (Figure S6E).

**Figure 6:**
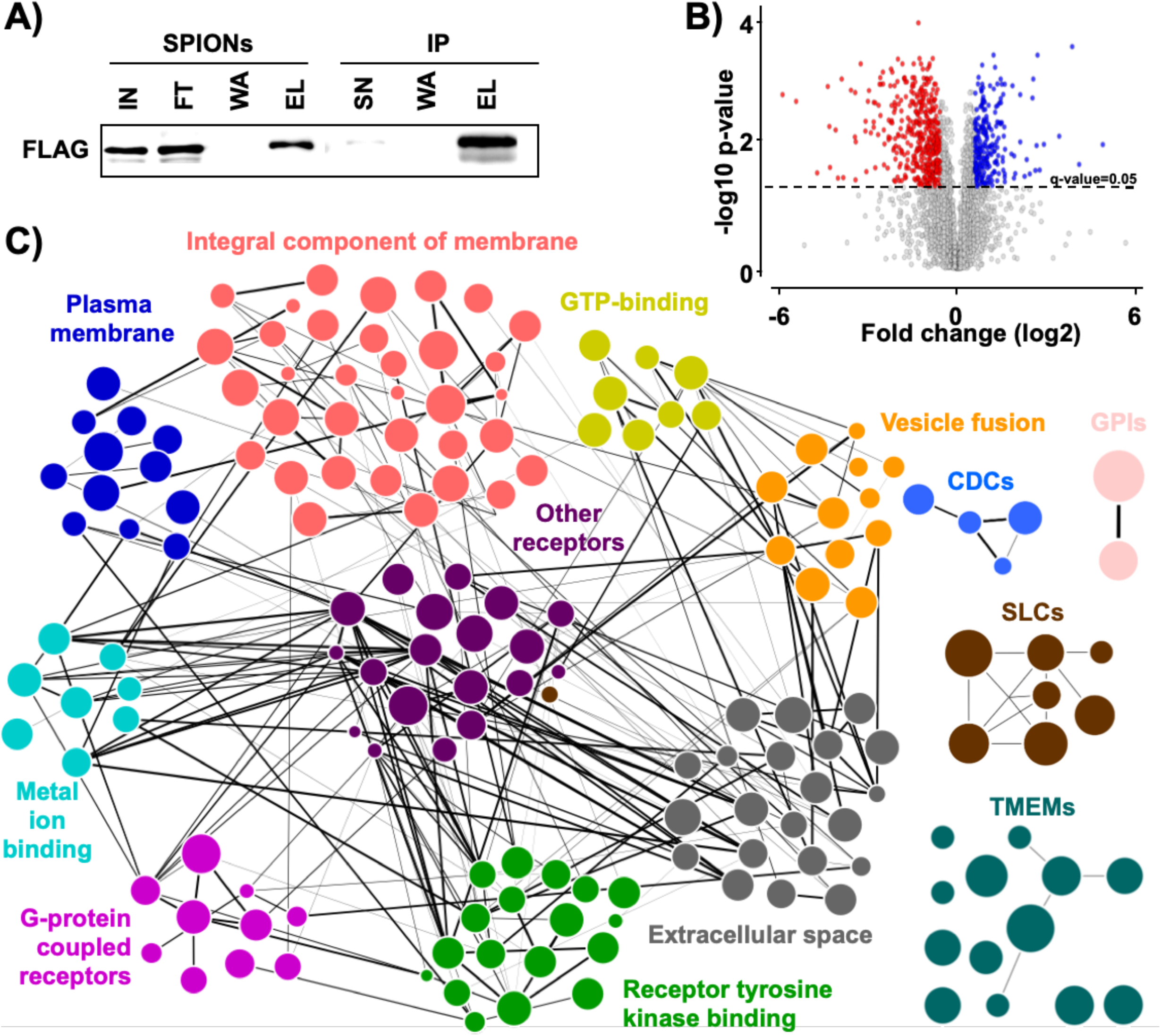
Identification of potential cargo of FLOT1-/FLOT2-positive early endosomes. **A)** Western blot analysis of FLAG-FLOT1/FLOT2 in early endosome-enrichment (SPIONs) and intact endosome-IP fractions. **B)** Data dependent acquisition (DIA)-based protein abundance fold-change ratios of SPIONs/SPIONs+IP fractions (n=3). Significantly different proteins are indicated. Cut offs: q-value: 0.05, fold change: 1.5. **C)** STRING-based PPI analysis of proteins overrepresented in SPIONs+IP fraction. Node size corresponds to DIA signal intensity and line thickness to PPI confidence score. **Abbreviations:** SPIONs: superparamagnetic iron oxide nanoparticles; IP: immunoprecipitation; IN: input; SN: supernatant; FT: flow through; WA: wash; EL: eluate; PPI: protein-protein interaction. **See also:** Figure S6.

The enriched population, which presents potential cargo of FLOT1-/FLOT2-positive early endosomes, consists of 328 proteins. GO-enrichment analysis revealed an overrepresentation of membrane proteins (Figure S6F), which is in agreement with a potential role of flotillins in the endocytosis and vesicular transport of plasma membrane proteins, demonstrated previously e.g. for NPC1L1 (Ge et al., 2011; Meister and Tikkanen, 2014). We subsequently performed STRING analyses to sub-classify the potential cargo proteins (Figure 6C). They contained both receptor tyrosine kinases and G-protein coupled receptors (50 receptors total), seven members of the solute carrier (SLC) family of transporters, and 13 members of the transmembrane protein (TMEM) family.

One group of receptors that sparked our interest was the latrophilins, as we detected all three members of this G protein-coupled receptor subfamily (LPHN1, LPHN2, and LPHN3) to be significantly enriched in FLOT1-/FLOT2-positive endosomes (Figure 7A). We confirmed their co-localization to FLOT1-/FLOT2-positive endosomes by immunofluorescence in HeLa and HEK293 cells (Figure 7 B, Figure S7). To further investigate the relation between FLOT1/FLOT2 and LPHN1/LPHN2/LPHN3, we performed IP experiments. We were able to co-IP all latrophilins with both flotillins, implicating not only the presence of latrophilins at FLOT1-/FLOT2-positive endosomes, but also a direct interaction (Figure 7C).

**Figure 7:**
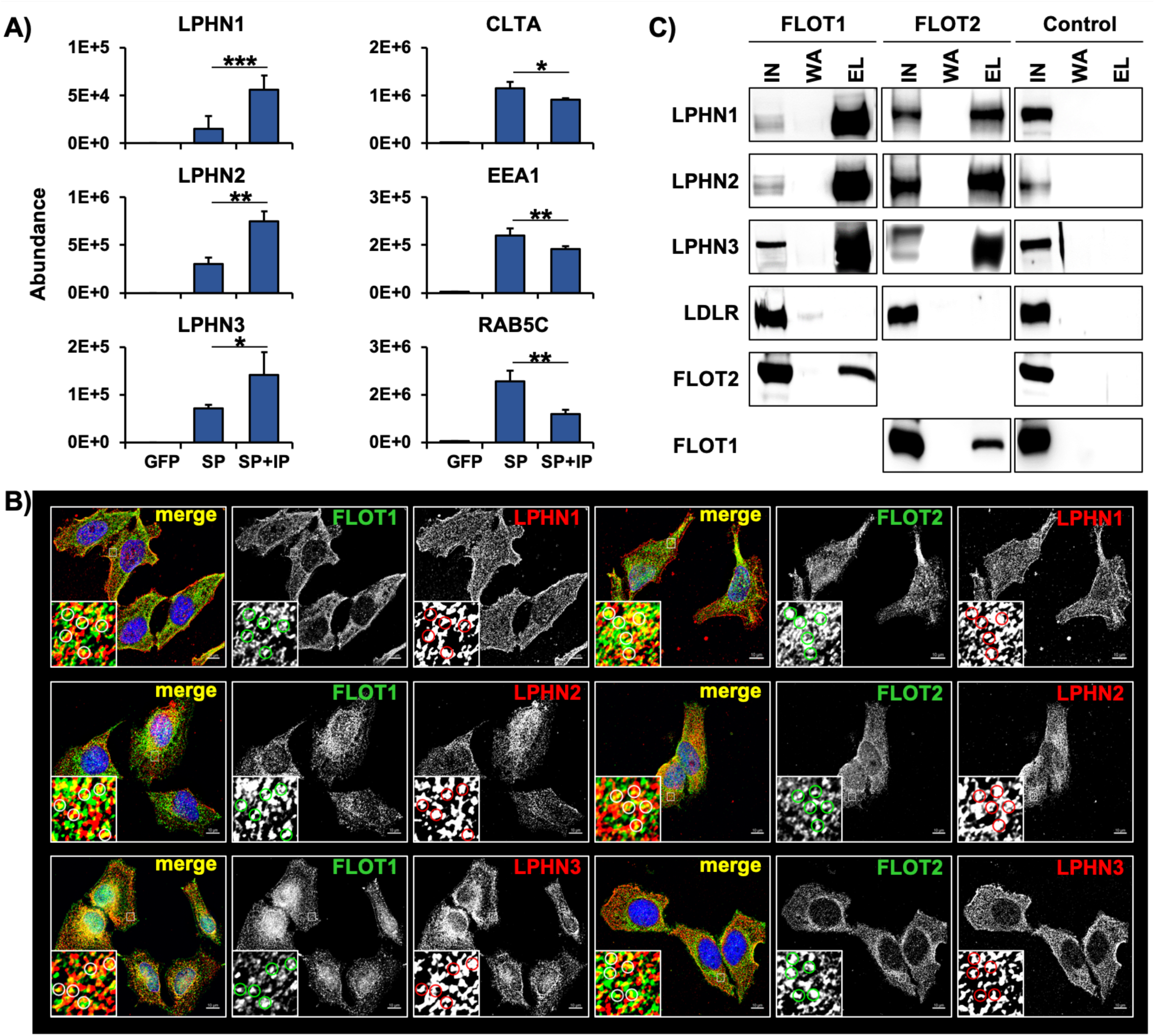
Latrophilins are a putative cargo of FLOT1-/FLOT2-positive early endosomes. **A)** Average DIA abundances for LPHN1, LPHN2, LPHN3, CLTA, EEA1, and RAB5C (n=3, +STDEV). Significance based on Student’s unpaired two-sided t-test: ∗ (p < 0.05), ∗∗ (p < 0.01), ∗∗∗ (p < 0.001). **B)** Co-immunostaining of HeLa cells for FLOT1, FLOT2, LPHN1, LPHN2, and LPHN3. **C)** Co-IP of FLOT1 and FLOT2 with LPHN1, LPHN2, and LPHN3. LDLR serves as a control for plasma membrane and clathrin-mediated endocytosis. **Abbreviations:** SP (SPIONs): superparamagnetic iron oxide nanoparticles; IP: immunoprecipitation; IN: input; WA: wash; EL: eluate. **See also:** Figure S7.

## DISCUSSION

In the current study, we present the first XL-LC-MS/MS analysis of lysosome-enriched fractions. In comparison to whole-cell studies, we detected significantly higher numbers of cross-links for proteins reported previously to be related to the lysosome. This was especially the case for *bona fide* lysosomal proteins, which are localized in/at the organelle (Thelen et al., 2017). When considering this shortlist, we identified >100-fold more cross-links compared to previously published whole-cell studies (Chavez et al., 2019; Liu et al., 2015). A probable key factor in this context is our starting material, consisting of SPIONs-enriched lysosomes, as, relative to the total proteome, lysosomal proteins are typically of low abundance (estimated 0.2 % of cellular protein mass (Itzhak et al., 2016)). With respect to individual lysosomal proteins, we did not observe a strong correlation of their abundance and the number of cross-links identified. We identified, for example, no cross-links for the lysosome-associated membrane glycoproteins 1/2 (LAMP1 and LAMP2), which were estimated to contribute 50 % of lysosomal protein mass (Eskelinen et al., 2004) and were among the most abundant lysosomal proteins in our dataset. Also, for the lysosomal luminal hydrolase CTSD, which was detected with the third-highest iBAQ value in the whole dataset, only two unique cross-links were detected, while we found the same number for the low abundant protein CHID1 (∼275-fold less abundant).

Based on the identified cross-links, we generated a protein interaction network containing 70 % potential novel PPIs. Compared to other approaches, cross-linking allows identifying interacting residues between individual proteins *in situ*. An approach able to generate data under similar near-native conditions is proximity biotinylation, utilizing e.g., BirA* or APEX2, which have been used in two different studies to investigate the lysosomal (surface) proteome and interactome *in viv*o (Go et al., 2021; Liao et al., 2019) In contrast to cross-linking, proximity biotinylation only allows to determine the presence of a protein within a defined radius relative to the respective fusion construct. It cannot identify, however, if two proteins are directly interacting, or which residues/domains are close to each other. This is exemplified by a recent study utilizing LAMP1, LAMP2, and LAMP3 fused to BirA* for investigation of the lysosomal proteome (Go et al., 2021) Comparison of the interaction partners identified for all three constructs revealed that only 8-18 % were unique, while the others were enriched in at least two datasets. Therefore, the interactome presented in this study provides a level of detail that is unprecedented for the analysis of lysosomal PPIs.

Another common approach for the investigation of protein interactions is co-IP, which we used to investigate two PPIs of members of the largest interaction networks, namely the FLOT1/FLOT2 complex and the V-ATPase complex. We selected the interaction of ATP6V1D with FZD9, a G-protein coupled multi-pass transmembrane receptor for WNT2 (Karasawa et al., 2002), as another member of this family, FZD8, was previously identified to interact with ATP6AP2. Intriguingly, it was shown in this study that correct V-ATPase function, and accordingly acidification of the lysosome, is necessary for WNT signaling, and that interaction of FZD8 with the V-ATPase complex plays a decisive role (Hermle et al., 2010). The direct interaction of ATP6V1D and FZD9 identified in this study substantiates this functional connection of WNT signaling and lysosomal acidification. A possible role could be related to lysosomal acidification. V-ATPase can only acidify lysosomes when the V0 part, which is integrated in the lysosomal membrane, pairs with the cytosolic V1 part. The independently assembled V1 part (Wang, 2020) can reversibly dissociate from V0, and this appears to be a process that can be regulated through different types of stimuli (Maxson and Grinstein, 2014). Similarly, it is conceivable that frizzled proteins could control V1/V0 V-ATPase assembly, and thus possibly regulate lysosomal acidification, and hence WNT signaling.

Another major lysosomal complex covered in our dataset was mTORC1. Among others, we detected cross-links for the interaction of the same lysine in the N-terminal region of LAMTOR1 with different residues of LAMTOR3. Both proteins are members of the Ragulator complex, whose interaction with the RAG-GTPases regulates mTORC1 activity (Zhang et al., 2017). It was shown previously that the N-terminal region of LAMTOR1 could not be crystallized without the RAG GTPases, implicating that it could exist in an unordered state under these circumstances (De Araujo, 2017). Matching of the three cross-links to the Ragulator crystal structure determined in the presence of RAG GTPases fulfilled only for one of them DSSO’s distant constraints. A possible explanation for the other two cross-links is that they originate from a state where Ragulator was not interacting with the RAG GTPases, indicating an alternative structure of the LAMTOR1 N-terminal region *in situ*.

The observation of several overlength cross-links, when matched to a structure determined from overexpression or *in vitro* experiments, can indicate that it does not present the native form of a protein (or it’s complex). In our dataset, the inter-links matched to the dimeric model of PPT1 (PDB: 3GRO) were responsible for 43 % of overlength cross-links, while all intra-links matched the published structure. Importantly, our proposed tetrameric PPT1 model satisfied all distance constraints. This is in line with previous findings, detecting the majority of enzymatic activity in size exclusion chromatography fractions correlating with the molecular weight of a tetrameric assembly (Lyly et al., 2007). The observed discrepancy to the crystallography-based structure could be related to the expression system utilized (*Spodoptera frugiperda*), as it lacks the capability to glycosylate PPT1 properly. It was shown previously, that glycosylation-deficient PPT1 variants are devoid of enzymatic activity, which was attributed to improper folding (Bellizzi et al., 2000; Lyly et al., 2007).

The flotillins, which yielded the highest number of cross-links, were reported in several studies to interact with a large variety of proteins at different subcellular locations, and to be involved in a plethora of processes (Bodin et al., 2014). FLOT1 has been demonstrated to play a role in clathrin/caveolin-independent endocytosis (Glebov et al., 2006), and the cargo of such FLOT1-positive endosomes has been shown to be delivered to lysosomes (Fan et al., 2019; Stuermer et al., 2001). Furthermore, FLOT1 was detected at the lysosomes’ cytosolic face (Kaushik, Kokuba). In accordance with these findings, we detected and verified the interaction of FLOT1/FLOT2 with guanine-nucleotide-binding protein subunit beta-4 (GNB4), a beta subunit of heterotrimeric G-proteins (Ruiz-Velasco et al., 2002). Variants of GNB4 were shown to cause Charcot–Marie–Tooth disease (Soong et al., 2013), and downregulation of GNB4 levels has been found in Gaucher Disease, a lysosomal storage disorder (Pawlinski et al., 2021) Association of GNB4 with lysosomes was implicated in a previous study (Schroder et al., 2007), and a yeast two-hybrid screen revealed its direct interaction with LAMP2 (Haenig et al., 2020) which co-localizes with FLOT1 and the lysosomal surface (Kaushik). Taken together, these data provide strong evidence that the lysosomal localization of GNB4 is mediated through interaction with the flotillins. With respect to the structure of flotillins, only rudimentary information existed based on NMR analysis of the mouse Flot2 N-terminal region (PDB: 1WIN). For the remaining protein, until recently, the predicted structure featured a 180° turn in the central helix of both flotillins resulting in direct interaction of the proteins’ N- and C-terminal domains (Rivera-Milla et al., 2006). The recently predicted FLOT1 and FLOT2 AlphaFold structures disagree with this model, predicting an extended helix with a bend in its middle for both of the proteins. This is supported by the cross-links detected in our dataset. We did not observe long-distance intra-links, which would be indicative for the proximity of distant regions of the proteins, and the inter-links of both proteins behave in unison, confirming intermolecular interactions along an extended region of the heterodimer.

We detected interaction with GNB4, among others, in the putative interaction hotspot, which localized in both flotillins to the major bend in the extended alpha-helical structure. Intriguingly, both the FLOT1 and the FLOT2 tyrosine residue located in this structure were reported in PhosphoSitePlus (Hornbeck et al., 2015) to be phosphorylated in >30 studies. Therefore, they could present a potential regulatory element for PPIs, e.g., to discriminate interactions regulated by tyrosine kinase- or G-protein-coupled-receptor signaling, as it was shown previously that flotillins are involved in both types of signaling pathways (Sugawara et al., 2007)

Multiple heptad coiled-coil motifs in both flotillins are known to favor higher-order protein structures (Grigoryan and Keating, 2008; Woolfson et al., 2012). In agreement with these characteristics and previously published results (Babuke et al., 2009; Solis et al., 2007), we identified different levels of higher-order flotillin structures, ranging from an abundant tetrameric assembly to assemblies exceeding 1MDa. Motivated by these findings, we generated a multi-heterodimeric model for the high molecular weight flotillin complex. Based on the close interaction of the FLOT1/FLOT2 PDZ3 binding motif, which is known to play a role in the assembly of large multiprotein complexes in stomatins (Hung and Sheng, 2002), and the reported 39-mer higher-order structure of rat liver vault protein (Tanaka, 2009), which shares characteristic features with the flotillins, we propose a model for a multimeric FLOT1/FLOT2 complex. A potential function for such an assembly could be the formation or transport of flotillin-positive endosomes (Glebov et al., 2006).

Except for a few proteins (Meister and Tikkanen, 2014), no substrates for flotillin-mediated endocytosis are known to date. Our analysis of its putative cargo identified >300 proteins, including a large number of membrane proteins and receptors. Among those, the latrophilins stood out, as all three members of this family of adhesion G protein-coupled receptors (Langenhan et al., 2016) were found to be significantly enriched. Furthermore, we observed both their co-localization and interaction with both flotillins, providing evidence for direct interaction, possibly in the course of flotillin-mediated endocytosis.

## METHOD DETAILS

### Cell culture and enrichment of lysosomes

Tissue culture plates (10 cm) were coated with 0.5 mg/mL poly-L-lysine (PLL) in 1x phosphate-buffered saline (PBS) for 20 min at 37 °C. On each plate, 6 x 10^6^ HEK293 cells were seeded in full medium supplemented with 10 % (v/v) superparamagnetic iron oxide nanoparticles (SPIONs) solution (DexoMAG40) and incubated for 24 h. Subsequently, cells were washed three times with pre-warmed PBS, fresh full medium was added, and cells were incubated for 24 h. Prior to harvesting, cells were washed twice with ice-cold PBS, scraped off the plates in 2 mL each of ice-cold isolation buffer (250 mM sucrose, 10 mM HEPES-NaOH pH 7.4, 1 mM CaCl2, 1 mM MgCl2, 1.5 mM MgAc, 1x cOmplete EDTA-free protease-inhibitor cocktail), and pooled. Cell suspensions from four plates each were dounced with 25 strokes in a 15 mL dounce homogenizer, and nuclei as well as intact cells were pelleted by centrifugation for 10 min at 600 x g, 4 °C. The supernatant (post-nuclear supernatant, PNS) was transferred to a new tube, the pellet resuspended in 3 mL of isolation buffer, and dounced and centrifuged again. The supernatant from this step was combined with the first one, and the pooled PNS was used for lysosome enrichment using LS columns in combination with a QuadroMACS magnet. Columns were equilibrated with 1 mL 0.5 % (w/v) bovine serum albumin (BSA) in PBS, the combined PNS of two cell culture plates was applied to one column, and the flow through was collected. After three washing steps with 1 mL isolation buffer each, columns were removed from the magnet, and lysosomes were eluted twice in 1 mL of isolation buffer using a plunger. Individual eluate fractions were centrifuged for 30 min at 20,000 x g, 4 °C, the supernatants were discarded, the pellets were resuspended in isolation buffer, and for each biological replicate, the pellets from 64 plates were pooled. Protein concentrations were determined using the DC protein assay. The efficiency of lysosome enrichment and lysosomal integrity was assessed using the β-hexosaminidase assay (Wendeler, 2009). Fractions obtained from lysosome enrichment (25 µL each) were combined with 8 µL of 10 % Triton X-100 or 8 µL of PBS, followed by the addition of 50 µL reaction solution (100 mM sodium citrate pH 4.6, 0.2 % (w/v) BSA, 10 mM para-nitrophenyl-N-acetyl-2-B-D-glucosaminide) in a 96 well plate format. Subsequently, the plate was incubated for 15 min at 37 °C, and 200 µL of stop solution (0.4 M glycine-HCl, pH 10.4) was added to the sample. Absorbance was measured at 405 nm on a microplate reader.

### Transfection of cells and enrichment of FLOT1-/FLOT2-positive early endosomes

64 tissue culture plates were coated with PLL (0.5 mg/mL), HEK293 cells were seeded at a density of 3.5 x 10^6^ cells/plate, and cultivated as described above. After 24 h, cells were transfected with 6 μg of plasmid (1:1 mixture of FLOT1-FLAG and FLOT2-FLAG or GFP) using TurboFect. After 4 h, the cell culture medium was replaced, and cells were incubated for 48 h. For enrichment of endosomes, SPIONs solution was added to the cells (10 % (v/v) final concentration) for a pulse period of 5 min. Subsequently, cells were washed with pre-warmed PBS and fresh full medium was added. Following a 5 min chase, the plates were placed on ice and endosomes were enriched following the same procedure as for the enrichment of lysosomes (see above). For IP of FLOT1-FLAG/FLOT2-FLAG-positive endosomes, obtained eluate fractions (1 mL each) were subsequently incubated with magnetic anti-FLAG beads (80 µL) on an end-over-end rotator at 4°C for 4 h. Beads were separated from samples by magnetic force (DynaMag 2 magnet), supernatants were transferred to new tubes, and the beads were washed three times with 500 µL PBS. For elution, beads were incubated with 150 µL of 150 ng/µL 3x FLAG peptide in 1x TRIS (hydroxymethyl) aminomethane buffered-saline (TBS, pH 7.6) for 30 min at 600 x g, 4 °C in a thermomixer. Subsequently, beads were separated from samples by magnetic force and eluate fractions transferred to a new tube. Protein concentrations were determined using the DC protein assay.

### Cross-linking of samples

For lysosome- and early endosome-enriched fractions, a portion of the eluate containing 500 µg and 200 µg protein, respectively, was transferred to a new tube. Intact organelles were pelleted by centrifugation for 30 min at 20,000 x g, 4 °C, the supernatant discarded, and the pellets resuspended in isolation buffer at a protein concentration of 1 mg/mL. Lysosomal samples were cross-linked in two states (intact and disrupted) while endosomal samples were cross-linked only in the intact state. For disruption of lysosomes, resuspended samples were lyzed with a sonicator (Bioruptor Plus) at an amplitude of 40 with 3 cycles of 30 sec each. All samples were cross-linked at final DSSO concentrations of 5, 2, 1, 0.5, 0.25 mM for investigation of the optimal cross-linker concentration (DSSO titration). For the XL-LC-MS/MS experiments of both lysosome- as well as endosome-enriched fractions, a final concentration of 0.25 mM DSSO was applied. After addition of DSSO, the cross-link reaction was allowed to proceed for 30 min at room temperature, and quenched by the addition of TRIS-HCl pH 8.0 (20 mM final concentration). Subsequently, proteins were precipitated by addition of acetone at a ratio of 4:1 (v/v) and incubation overnight at -20 °C. The next day, samples were centrifuged for 20 min at 20,000 x g, 4 °C, the supernatant was discarded, the pellet washed twice with ice-cold acetone, air-dried, and stored at -80 °C until further use.

### Sodium dodecyl sulfate-polyacrylamide gel electrophoresis (SDS-PAGE)

Polyacrylamide gels were prepared in-house. Both running and stacking gels were prepared with 10 % (w/v) SDS, 40 % (v/v) acrylamide, 10 % (w/v) ammonium persulfate (APS), and 1 % (v/v) tetramethylethylenediamine (TEMED), while 1.5 M TRIS-HCl pH 8.8 and 0.5 M TRIS-HCl pH 6.8 was used for running and stacking gels, respectively. Laemmli buffer (Laemmli, 1970) (4x stock, 240 mM TRIS-HCl pH 7.4, 4 % (v/v) β-mercaptoethanol, 8 % (w/v) SDS, 40 % (v/v) glycerol, 4 % (w/v) bromophenol blue) was added to samples (1x final concentration) followed by incubation for 10 min at 56 °C. Gel electrophoresis was performed at 80-140 V for up to 1.5 h. Gels were either stained overnight with Coomassie Brilliant Blue G-250 or with a Silver Stain Kit.

### Immunoprecipitation of endogenous proteins

HEK293 cells were seeded at a density of 6 x 10^6^ cells/10 cm plate and cultivated for 48 h in full medium. Cells were washed twice with ice-cold PBS, scraped off the plate in 1 mL of ice-cold PBS, transferred to a microtube, and centrifuged for 10 min at 600 x g, 4 °C. The supernatant was discarded and the pellet resuspended in 300 µL of RIPA lysis buffer (50 mM TRIS-HCl pH 7.4, 150 mM NaCl, 1 % Triton X-100, 0.1 % (w/v) SDS, 0.5 % (w/v) sodium deoxycholate (SDC), 1x cOmplete EDTA-free protease-inhibitor cocktail, 1 mM EDTA). Samples were incubated on ice for 30 min and passed through a 25 Gauge needle every 10 min. Subsequently, the lysate was cleared by centrifugation for 15 min at 20,000 x g, 4 °C, transferred to a new pre-cooled microtube, and protein concentrations were determined using the DC protein assay. For each sample, lysate containing 1.6 mg of protein was incubated with 3 µg of antibody overnight by end-over-end rotation at 4 °C. The next morning, 60 µL of Protein A beads were added to each sample, followed by end-over-end incubation for 1 h at 4 °C. Beads were pelleted by centrifugation for 5 min at 1000 x g, 4 °C, supernatants transferred to new tubes, and beads were washed three times with 500 µL of ice-cold PBS. Proteins were eluted from the beads by incubation in 2x Laemmli buffer for 30 min at 45 °C.

### Blue native polyacrylamide gel electrophoresis (BN-PAGE)

Gradient gels (4-13 %) were prepared with a gradient mixer and overlaid with a stacking gel (Table 1). Samples were supplemented with solubilization buffer (10 mM HEPES-NaOH pH 7.4, 1 % (v/v) digitonin, 2 mM EDTA, 50 mM NaCl, 10 % (v/v) glycerol, 1 mM phenylmethylsulfonyl fluoride (PMSF)) and loading dye (10 mM Bis-TRIS-HCl pH 7.0, 0.5 % (w/v) Coomassie Brilliant Blue G-250, 50 mM ε-amino n-caproic acid), and loaded to the gel placed in a pre-cooled gel chamber. The anode buffer was 50 mM Bis-TRIS-HCL pH 7.0 and the cathode buffer 50 mM Tricine pH 7.0, 15 mM Bis-TRIS-HCl pH 7.0, 0.2 % (w/v) Coomassie Brilliant Blue G-250. The temperature of the gel chamber was maintained at 4 °C and electrophoresis was performed at 50 V for 20 h.

**Table 1.** BN-PAGE gel setup.

| Component | Separating gel |  | Stacking gel |
| --- | --- | --- | --- |
|  | 4 % | 13 % | 3,5 % |
| 3x gel buffer (55 mM Imidazole, 1.5 M 6-Amino n-caproic acid) [mL] | 2 | 2.5 | 1 |
| Acrylamide/bis-acrylamide AB-3 mix (49.5 % T, 3 % C) [mL] | 0.5 | 1.9 | 0.2 |
| Glycerol [g] | - | 1.5 | - |
| 10 % (w/v) APS [μL] | 33.3 | 37.5 | 25 |
| TEMED [μL] | 3.3 | 3.7 | 2.5 |
| Water [mL] | 3.5 | 1.5 | 1.7 |
| Total [mL] | 6 | 7.5 | 3 |

### Western blotting

Proteins were transferred to nitrocellulose or polyvinylidene fluoride (PVDF) membranes using a semi-dry or wet electro blotter for 1 h or 2 h at 200 mA/membrane. Membranes were blocked in 5 % non-fat dry milk in TBS containing 0.05 % (v/v) Tween 20 (TBS-T) for 1 h at RT followed by incubation with primary antibodies overnight at 4 °C (for dilutions of individual antibodies see key resource table). The next day, membranes were washed three times with TBS-T for 10 min at RT followed by incubation with secondary antibody (for dilutions of individual antibodies see key resource table) for 60 min at RT. Subsequently, membranes were washed three times for 10 min at RT with TBS-T, and the blots were developed. Protein expression signals were detected using the enhanced chemiluminescence (ECL) kit, visualized with the FUSION SOLO 4M system, and analyzed by the FusionCapt advance software.

### Immunofluorescence microscopy

HeLa and HEK293 cells were seeded in 12-well plates at a density of 2 x 10^4^ and 40 x 10^4^ cells per well, respectively. For HEK293 cells, glass coverslips were coated using PLL (0.5 mg/mL). Cells were cultured for 36 h after seeding. Cells were transfected with 1 µg of plasmid (1:1 mixture of FLOT1-FLAG and FLOT2-FLAG) using TurboFect. After 4 h, the medium was replaced with fresh full medium and cells were incubated for 48 h. For staining, cells were washed using PBS and fixed with ice-cold methanol at -20 °C for 20 min. After fixation, cells were washed twice with PBS and blocked with 2 % BSA in PBS for 1 h at RT. Blocked cells were stained with primary (for dilutions of individual antibodies see key resource table) overnight at 4 °C in a humid chamber. Subsequently, cells were washed three times with TBS for 5 min each and incubated with secondary antibodies (for dilutions of individual antibodies see key resource table) for 1 h at RT in the dark. Coverslips were washed three times with TBS for 5 min each, rinsed once with distilled water, and mounted on specimen slides using ROTI Mount FluorCare DAPI. Images were acquired using an Axiovert 200 M microscope equipped with an AxioCam705 camera. Images were prepared using the ZEN software.

### Mass spectrometry sample preparation

Precipitated proteins were resuspended in 100 µL of freshly prepared 8 M urea/100 mM triethylammonium bicarbonate (TEAB) and incubated for 45 min at 600 rpm, 37°C. Disulfide bridges were reduced with 4 mM DTT (final concentration) at 56 °C for 30 min, alkylated with 8 mM chloroacetamide (final concentration) at RT for 30 min (Müller and Winter, 2017), and the reaction was quenched by the addition of 4 mM DTT. Subsequently, samples were diluted 1:1 with 100 mM TEAB, rLysC was added at an enzyme-to-protein ratio of 1:100 (w/w), and proteolytic digestion was performed at 37 °C overnight. The following day, the urea concentration was reduced to 1.6 M by addition of 100 mM TEAB, trypsin was added at an enzyme-to-protein ratio of 1:100 (w/w), and the samples were incubated at 37 °C for 8 h. The resulting peptides were desalted using 50 mg Sep-Pak C18 cartridges, dried using a vacuum centrifuge, and stored at -80 °C until further use.

### Strong cation-exchange (SCX) chromatography fractionation

SCX fractionation was performed with an UltiMate 3000 RSLC HPLC chromatography system in combination with a PolySULFOETHYL A column (150 mm x 1 mm, 5 µm particle size). Desalted peptides (500 µg each) were reconstituted in 20 µL of SCX solvent A (20 % acetonitrile (ACN), 10 mM KH_2_PO_4_ pH 2.7), loaded to the analytical column with 100 % SCX solvent A, and eluted with increasing amounts of SCX solvent B (500 mM KCl, 20 % ACN, 10 mM KH_2_PO_4_ pH 2.7) at a flow rate of 50 µL/min. The gradient was as follows (adapted from (Klykov et al., 2018)) 0-42 min: 0-2 % B; 42-50 min: 2-3 % B; 50-60 min: 3-8 % B; 60-70 min: 8-20 % B; 70-80 min: 20-40 % B; 80-86 min: 40-90 % B; 86-90 min: 90 % B; 90-91 min: 0 % B; 91-120 min: 0 % B. Eluting peptides were collected with a fraction collector, individual fractions dried using a vacuum centrifuge, resuspended in 20 µL of 5 % ACN, 5 % formic acid (FA), and desalted using C18 STAGE-tips (Rappsilber et al., 2007). STAGE tip eluates were dried using a vacuum centrifuge, resuspended in 5 % ACN, 5 % FA, and peptide concentrations were determined with a quantitative fluorometric peptide assay.

### Liquid chromatography electrospray ionization tandem mass spectrometry (LC-ESI-MS/MS)

Dried peptides were resuspended in 5 % ACN, 5 % FA and analyzed using an UltiMate 3000 RSLCnano UHPLC system coupled to an Orbitrap Fusion Lumos mass spectrometer. From each sample (25 % of the total amount for individual SCX fractions, 1 µg of non-cross-linked samples) were loaded on a 50 cm C18 reversed-phase analytical column at a flow rate of 600 nL/min using 100 % solvent A (0.1 % FA in water). For analytical column preparation, fused silica capillaries (360 μm outer diameter, 100 μm inner diameter) were used to generate spray tips using a P-2000 laser puller. Tips were packed with 1.9 μm ReproSil-Pur AQ C18 particles to a length of 50 cm. Peptide separation was performed with 120 min (SCX fractions) and 240 min (non-cross-linked samples) linear gradients from 5-35 % solvent B (90 % ACN, 0.1 % FA) at a flow rate of 300 nL/min. MS1 spectra were acquired in the Orbitrap mass analyzer from m/z 375-1,575 at a resolution of 60,000. For peptide fragmentation, charge states from 3+ to 8+ for cross-linked samples and 2+ to 5+ for non-cross-linked samples were selected, and dynamic exclusion was defined as 60 sec and 120 sec for 120 min and 240 min gradients, respectively.

Cross-linked samples were either analyzed with an MS2-MS3-MS2 strategy (Liu et al., 2017) or with a stepped collision energy approach (Stieger, 2019), where ions with the highest charge state were prioritized for fragmentation. For both methods, MS2 scans were acquired at a resolution of 30,000 in the Orbitrap analyzer with a dynamic mass window. In case of stepped collision energy, peptides were fragmented using higher collision dissociation (HCD) with 21, 26, and 31 % normalized collision energy (NCE). For the MS2-MS3-MS2 fragmentation method, sequential collision-induced dissociation (CID) and electron-transfer dissociation (ETD) spectra were acquired for each precursor. The precursor isolation width was set to m/z 1.6 with standard automatic gain control and automatic maximum injection time. The NCE for CID-MS2 scans were set to 25 % and calibrated charge-dependent ETD parameters enabled. MS3 scans were triggered by a targeted mass difference of 31.9721 detected in the MS2 scan. The MS3 scan was performed in the ion trap part of the instrument with CID at 35 % NCE with a normalized automatic gain control (AGC) target of 300 %.

Data independent acquisition (DIA) analyses were performed with the same instrumental setup as described above. For each sample, 1 µg of peptides were loaded directly on a reversed-phase analytical column packed with 3 μm ReprosilPur AQ C18 particles to a length of 40 cm and separated with 120 min linear gradients. After acquisition of one MS1 scan 24 static window DIA MS2 scans were performed. MS1 scans were acquired in the Orbitrap analyzer from m/z 350-1200 at a resolution of 120,000 with a maximum injection time of 20 msec and an AGC target setting of 5 x10^5^. MS2 scans were defined to cover the MS1 scan range with 36 scan windows of 24.1 m/z each, resulting in an overlap of 0.5 m/z and a cycle time of 3.44 sec. Peptides were fragmented by HCD with an NCE of 27 %, and spectra were acquired in the Orbitrap analyzer with a resolution of 30,000, a maximum injection time of 60 msec, and an AGC target setting of 1x10^6^.

### Proteome Discoverer analysis

Thermo *.raw files from cross-linked samples were analyzed using Proteome Discoverer, utilizing Mascot and XlinkX for peptide identification. The following settings were used for both algorithms: precursor ion mass tolerance: 10 ppm; Orbitrap fragment ion mass tolerance: 20 ppm; ion trap fragment ion mass tolerance: 0.5 Da; fixed modification: carbamidomethylation at cysteine; variable modification: oxidation at methionine; enzyme: trypsin; number of allowed missed cleavage sites: 2; minimum peptide length: 5 amino acids; cross-linking site: lysine (K) and N-terminus of proteins. Data were searched against UniProt *Homo sapiens* (Entries: 20,365) in combination with the common repository of adventitious proteins (cRAP) database containing common contaminants. The Proteome Discoverer workflow was split into two branches with a cross-link and standard peptide search. MS2 spectra containing DSSO reporter ions were analyzed with pre-defined “MS2-MS2-MS3” and “MS2” search options using XlinkX. Peptide identifications were accepted with a minimum XlinkX score of 40 and filtered at false discovery rates (FDRs) of 1 % and 5 % at the cross-linked peptide spectrum level. Cross-links were exported. Spectra, which did not contain reporter ions were searched using Mascot. Identified peptides were filtered at 1 % FDR on the peptide level using Percolator and proteins exported at 1 % FDR. Only high confidence peptide identifications were considered and data exported. Data from both algorithms were further analyzed applying different software packages (R, Excel, GraphPad Prism, STRING, Cytoscape, xiVIEW, TopoLink, PSIPRED, PCOILS, and PyMol)

### MaxQuant analysis

Thermo *.raw files from non-cross-linked samples were analyzed using MaxQuant (Cox and Mann, 2008) for determining iBAQ values (Schwanhausser et al., 2011). The following settings were used: precursor ion mass tolerance: 4.5 ppm; Orbitrap fragment ion mass tolerance: 20 ppm; fixed modification: carbamidomethylation at cysteine; variable modifications: oxidation at methionine, acetylation at protein N-terminus, and deamidation at asparagine (N) as well as glutamine (Q); enzyme: trypsin; number of allowed missed cleavage sites: 2; minimum peptide length: 5 amino acids. Data were searched against UniProt *Homo sapiens* (Entries: 20,365) in combination with the cRAP database containing common contaminants. Data was filtered at 1 % FDR on the peptide level and protein level and exported, followed by analysis with different software packages (Excel and GraphPad Prism).

### Spectronaut analysis

Thermo *.raw DIA files from FLOT1-FLAG+FLOT2-FLAG-transfected HEK293 cells were analyzed using Spectronaut. Initially, hybrid spectral libraries were generated from both DDA and DIA files with the Pulsar search engine integrated into Spectronaut, applying the following parameters: precursor ion mass tolerance: dynamic; Orbitrap fragment ion mass tolerance: dynamic; fixed modification: carbamidomethylation at cysteine; variable modifications: oxidation at methionine, acetylation at protein N-terminus and deamidation at asparagine (N) as well as glutamine (Q); enzyme: trypsin; number of allowed missed cleavage sites: 2; minimum peptide length: 5 amino acids. Data were searched against UniProt *Homo sapiens* (Entries: 20,365) in combination with the cRAP database containing common contaminants. For each peptide, the 3 - 6 most abundant b/y ions were selected for library generation, dependent on their signal intensity. Dynamic retention time alignment was performed based on the high-precision indexed retention time (iRT) concept (Bruderer, 2016). Mass tolerances (precursor and fragment ions) as well as peak extraction windows were defined automatically by Spectronaut. Normalization was disabled and data filtered at 1 % FDR on the peptide and protein level (q-value < 0.01). High confidence identifications were exported, followed by analysis with different software packages (R, Excel, STRING, Cytoscape, and GraphPad Prism).

### Structural analysis

Protein sequences were obtained from UniProt and searched using BLAST against protein database (PDB) entries with an E-value of 0.0001. In case no reference structure for *Homo sapiens* was available, structures from other organisms were obtained from the SWISS-MODEL repository (Waterhouse et al., 2018), and/or predicted structures from the AlphaFold protein structure database were used (Jumper et al., 2021). Amino acid numbering was adjusted to UniProt entries, identified cross-links were mapped and topologically evaluated with TopoLink (Ferrari et al., 2019), and visualized using PyMol.

### Molecular docking

Protein structure perturbation and optimization was performed with SCWRL (Krivov et al., 2009) and restraint-based docking with HADDOCK (Van Zundert et al., 2016) as well as CNS (Brunger, 2007). Distance constraints of identified cross-links (20 ± 10 Å) were used to limit the possible interaction search space, applying unambiguous restraint distances on C-beta (except for glycine, for which C-alpha was used). In line with the default HADDOCK protocol, 500 initial restraints-based complex models were generated, followed by their rigid-body energy minimization. For the best 100 models, semi-flexible refinement in torsion angle space was performed, followed by molecular dynamics refinement in explicit water. Generated models were evaluated based on the weighted sum of electrostatic and van der Waals energies, complemented by the empirical desolvation energy. Based on these parameters, the ten best-scoring models were reported. Finally, models were further clustered within a 5 Å pairwise root mean square deviation, and the lowest energy model of each cluster reported. Results were visualized using PyMol.

## QUANTIFICATION AND STATISTICAL ANALYSIS

Spectronaut results were analyzed using R (R Core Team, 2020) applying the integrated development environment RStudio. For data importing, tidying, transforming and statistical modeling Tidyverse (Wickham, 2019) and R base packages were used. Results were exported using Openxlsx and visualized by Ggplot and Viridis. Protein signal intensities were initially log2-transformed for data quality assessment and visualization. Missing values were replaced by "NA", while imputation of missing values was omitted. Subsequently, data from individual replicates of the experimental conditions (GFP, SPIONs, and SPIONs+IP) were categorized into three populations as follows: background (three valid values in all three conditions), SPIONs-specific (no valid GFP value and ≥ 2 valid values in SPIONs or SPIONs+IP) and SPIONs “on/off” (≥ 2 valid values in only one SPIONs condition and no valid value in other conditions). Subsequently, non-logarithmic data of the individual data sets were normalized on the signal intensities of FLOT1 and FLOT2 in the respective datasets, while the GFP sample was not normalized, followed by log2 transformation of all datasets. Proteins with ≥ 2 valid values in each dataset were compared using a two-sided unpaired t-test. On/off proteins were defined as significant and their p-values set to zero; while p-values of proteins not matching any of the two conditions were set to one. P-value-adjustment was performed according to Benjamini-Hochberg (Benjamini and Hochberg, 1995) and proteins with a q-value ≤ 0.05 were considered differently enriched. Significantly enriched proteins were submitted to protein-protein interaction and enrichment analysis with STRING (Szklarczyk et al., 2019) while the entire list of proteins found in the experiment was defined as background gene set. Networks (type: full) were generated, in which the edges indicate both functional and physical protein associations.

---

**Figure S1:**
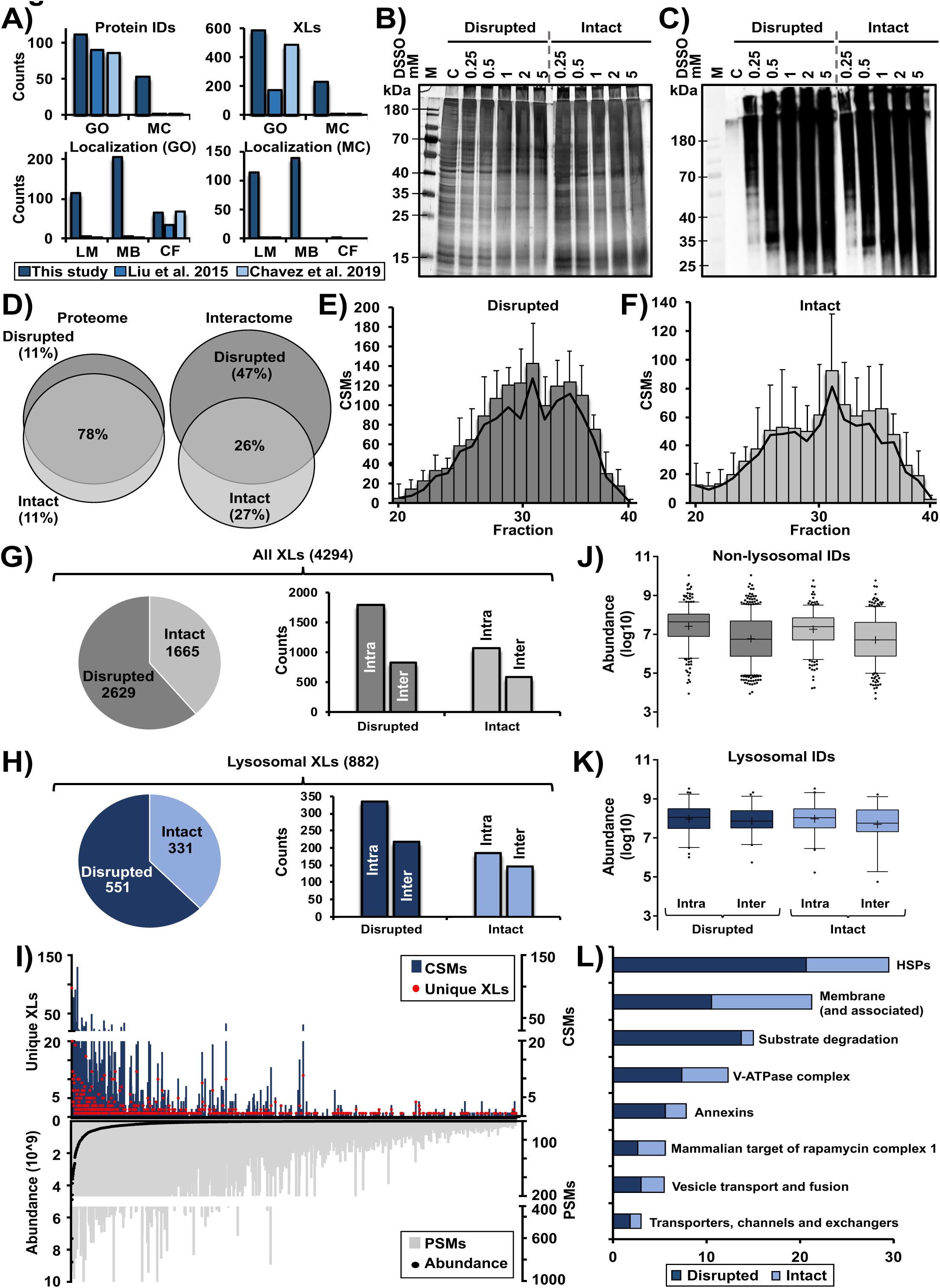
Cross-linking mass spectrometry analysis of HEK293 lysosome-enriched fractions. Related to Figure 1. **A)** Identified cross-links for lysosomal and lysosome-related proteins. Proteins are categorized in two groups: GO: proteins which are related to lysosomes based on their categorization in gene ontology databases, also including proteins which are not important for lysosomal function; MC manually curated list of proteins which are localized in/at lysosomes (Thelen et al., 2017) and known to play a role in lysosomal function. Numbers from the published whole proteome cross-linking studies were extracted from the respective supplementary tables. Cross-links assigned to both GO and MC were further subcategorized based on the protein’s location at the lysosome **B)** Silver-stained SDS-PAGE gel of lysosome-enriched fractions cross-linked with the indicated amounts of DSSO in a disrupted and intact state at a protein concentration of 1 µg/µL. For each sample, 2.5 µg of protein were loaded to the gel. Control samples were treated with DMSO. Based on the band pattern, the optimal concentration of DSSO was determined to be **I)** 0.25 mM (final concentration), as the application of higher amounts resulted in a shift of band patterns. Especially with respect to the lower molecular weight region, less bands were observed, while the high molecular weight region (containing cross-linked proteins) increased in intensity, which is indicative for protein aggregates resulting from over-cross-linking. Shown are two separate SDS-PAGE analyses (performed under the same conditions) for samples cross-linked in the intact and the disrupted state, individual gels are indicated by a dashed grey line. **C)** Western blot analysis of samples described in (B) for the visualization of cross-linker-modified proteins using an in-house produced antibody which shows immunoreactivity against water-quenched DSSO (Singh et al., 2021). For each sample, 1 µg of protein was loaded to the gel. Shown are two separate western blot analyses (performed under the same conditions) for samples cross-linked in the intact and the disrupted state, individual gels are indicated. **D)** Overlap in proteins, which are not categorized as lysosomal based on their GO term, identified in mass spectrometric analyses of lysosome-enriched fractions in the disrupted and the intact state. Proteome: proteins identified in analyses of non-cross-linked samples; Interactome: unique cross-links identified in analyses of cross-linked samples. **E/F)** Distribution of identified CSMs in LC-MS/MS analyses of SCX-fractionated tryptic digests of lysosome-enriched fractions cross-linked in a disrupted (E) and intact (F) state. Since later eluting SCX fractions contain the majority of higher charged cross-linked tryptic peptides, only SCX fractions 20-40 were analyzed by LC-MS/MS. Shown are average values (n=3, +STDEV). **G/H)** Distribution of identified cross-links from the analysis of lysosome-enriched fractions cross-linked in the disrupted and the intact state. Shown are numbers identified in the disrupted/intact state, and how many cross-links were assigned as intra- or inter-links, for cross-links identified in the whole dataset (G) and such assigned to lysosomal proteins (H). **I)** Correlation of cross-link identification and protein abundance for all non-lysosomal proteins. Proteins are sorted based on their average iBAQ abundance (Schwanhausser et al., 2011) in the LC-MS/MS analysis of non-cross-linked samples. Average iBAQ abundances as well as total numbers of unique cross-links, CSMs, and PSMs are shown. Values are based on the analysis of (n=3) each for lysosome-enriched fractions in the intact and disrupted state. **J/K)** Distribution of iBAQ abundances for all (J) and lysosomal proteins (K) for which intra- or inter-links were detected. Values are based on the LC-MS/MS analysis of non-cross-linked lysosome-enriched fractions in the disrupted and the intact state. Only proteins for which cross-links were identified in the respective state were considered. Shown are combined values from three independent replicates, the median is indicated by a line and the average by “+”. **L)** Categorization of the 1,415 CSMs identified for 68 lysosomal proteins. Proteins were assigned to previously defined categories of lysosomal proteins (Akter, 2020). **Abbreviations:** IDs: identifications; LM: lumen; MB: membrane; CF: cytosolic face; GO: gene ontology; MC: manually curated; DSSO: disuccinimidyl sulfoxide; DMSO: dimethyl sulfoxide; C: control (no DSSO); M: protein marker; iBAQ: intensity based absolute quantification; SCX: strong cation exchange; XL: cross-link; PSMs: peptide spectral matches; CSMs: cross-link spectral matches; HSPs: heat shock proteins.

**Figure S2:**
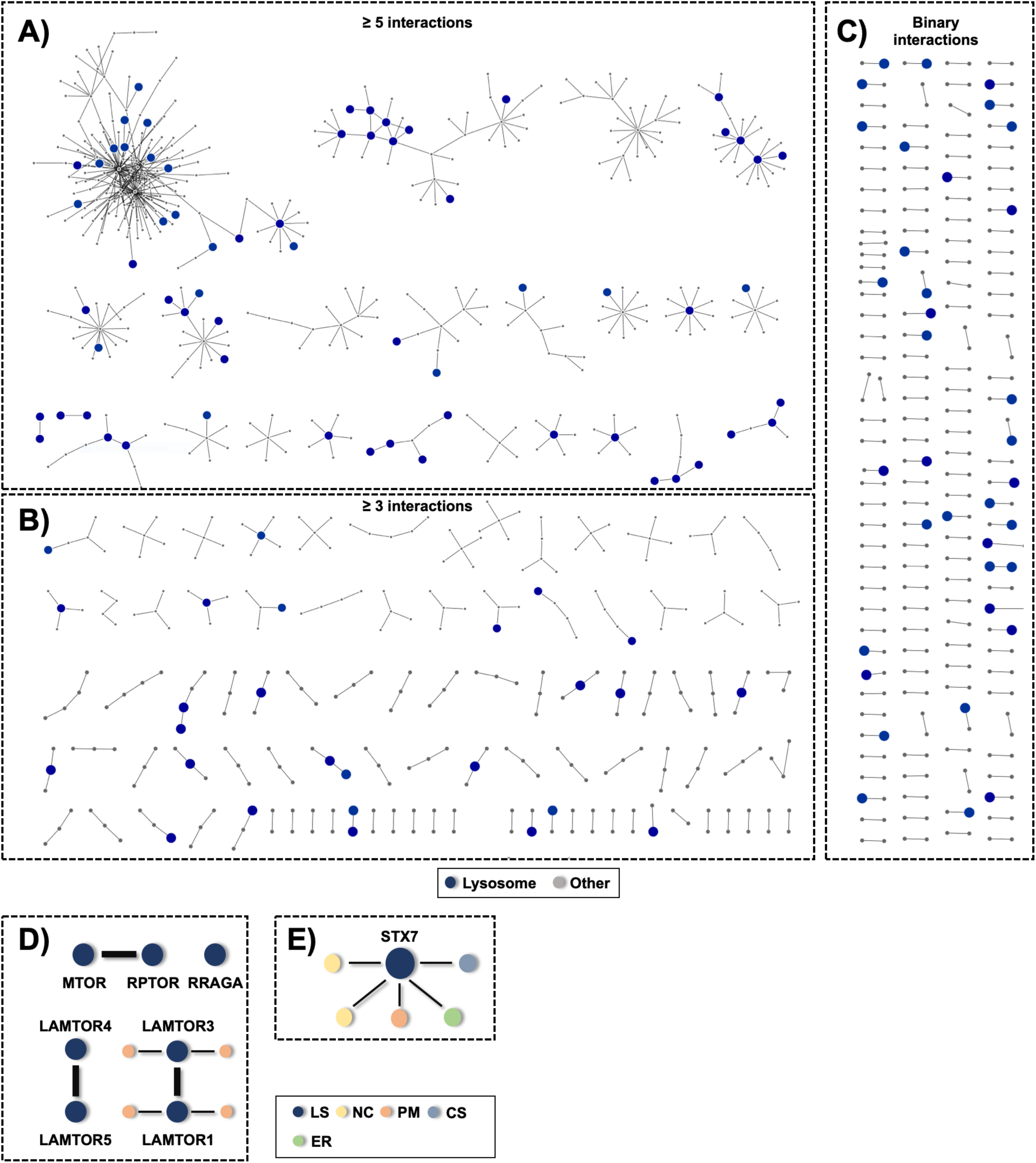
Analysis of protein-protein interactions in lysosome-enriched fractions. Related to Figure 2. **A-C)** Protein-protein interaction networks based on cross-links identified between two different proteins (inter-links, 1,023 in total). Lysosomal proteins are highlighted as large blue filled circles while non-lysosomal proteins are depicted as small grey dots. Interactions were extracted from the entire XL-LC-MS/MS dataset from disrupted and intact lysosomes. Based on the number of interactors involved in a particular network, data were grouped into different classes: A) ≥ 5 interactors, B) ≥3 interactors, C) binary interactions. The networks were generated using Cytoscape. **D)** Protein-protein interaction networks for proteins related to mTORC1; subcellular localization of interactors is indicated by color. The networks were generated using Cytoscape. **E)** Protein-protein interactions identified for Syntaxin 7 (STX7), subcellular localizations of interactors are indicated by color. The network was generated using Cytoscape. **Abbreviations:** LS: lysosome; NC: nucleus; PM: plasma membrane; CS: cytoskeleton; ER: endoplasmic reticulum.

**Figure S3:**
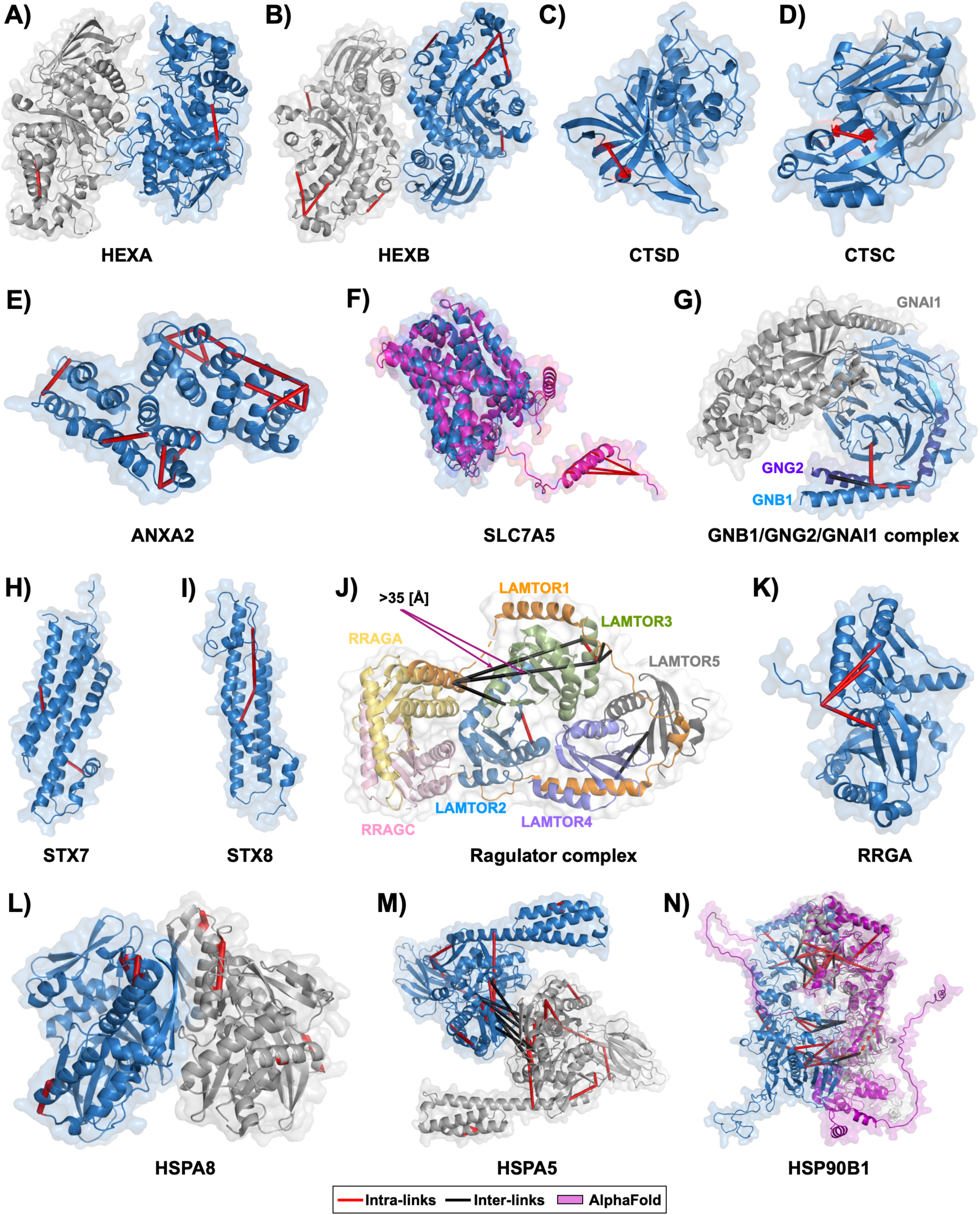
Integration of cross-links into resolved structures and AlphaFold models of lysosomal and lysosome-associated proteins. Related to Figure 3. **A)** Integration of cross-links into the RS of beta-hexosaminidase subunit alpha (HEXA). **B)** Integration of cross-links into the RS of Beta-hexosaminidase subunit beta (HEXB). **C)** Integration of cross-links into the RS of cathepsin D (CTSD). **D)** Integration of cross-links into the RS of cathepsin C (CTSC). **E)** Integration of cross-links into the RS of annexin A2 (ANXA2). **F)** Integration of cross-links into the AF (Jumper et al., 2021) model of the large neutral amino acids transporter small subunit 1 (SLC7A5). **G)** Integration of cross-links into the RS of the complex of GNB1, GNG2, and GNAI1. **H)** Integration of cross-links into the homology model of syntaxin 7 (STX7). **I)** Integration of cross-links into the homology model of syntaxin 8 (STX8). **J)** Integration of cross-links into the RS of the Ragulator complex (LAMTOR1-5) in the presence of RRAGA and RRAGC. **K)** Integration of cross-links into the AF model of Ras-related GTP-binding protein A (RRGA). **L)** Integration of cross-links into the RS of heatshock protein 8 (HSPA8). **M)** Integration of cross-links into the RS of heatshock protein 5 (HSPA5). **N)** Integration of cross-links into the mixed model based on the RS and the AF model for heatshock protein 90B1 (HSP90B1). Magenta structures were extracted from AlphaFold while grey/blue structures are based on crystal or cryo-EM structures retrieved from PDB. **Legend:** Inter-links (cross-links between two different proteins or two subunits of the same protein) are shown in black, and intra-links (cross-links within the same protein) are shown in red. Overlength cross-links (>35 Å) are indicated.

**Figure S4:**
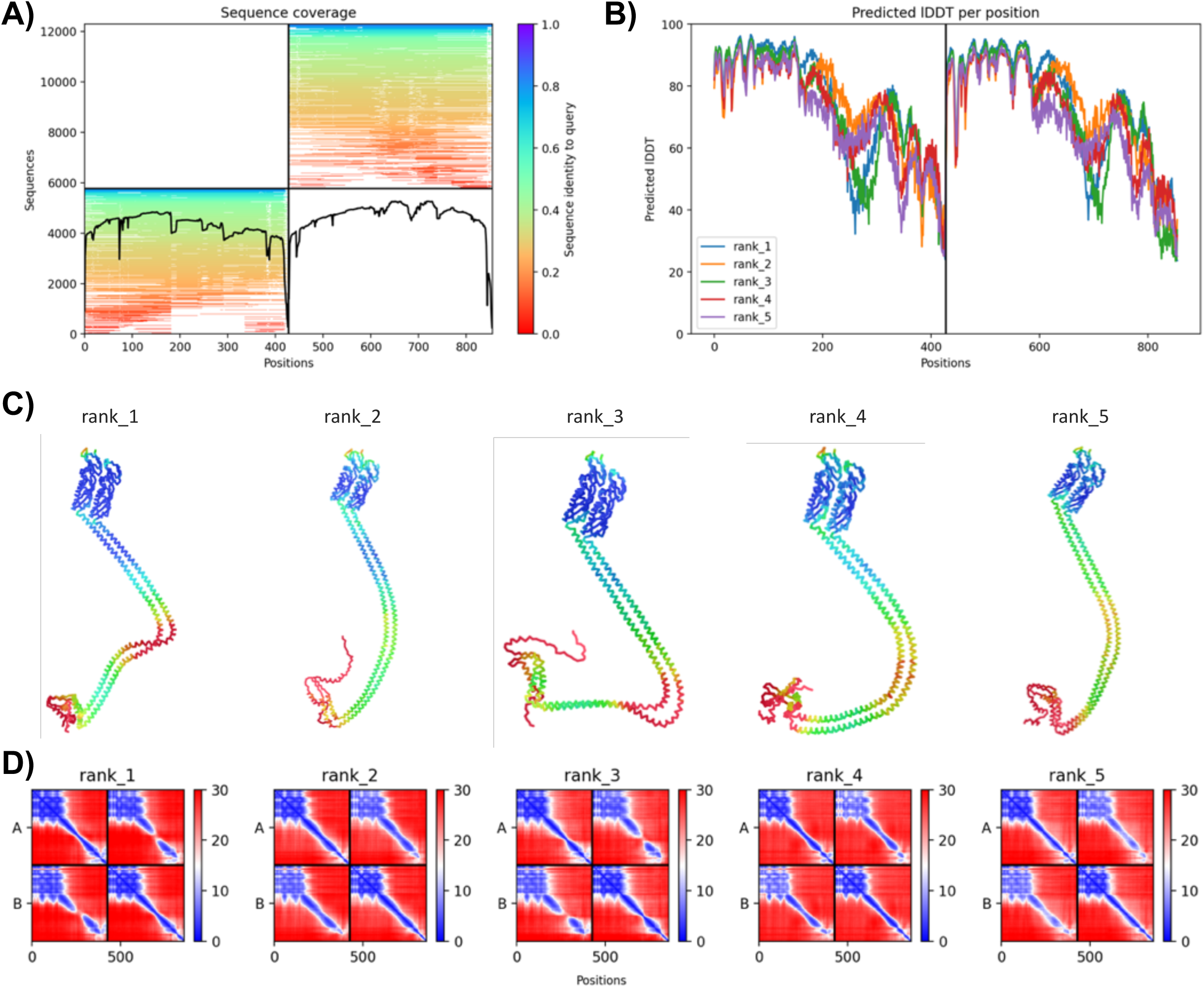
Quality assessment of different AlphaFold-based ColabFold models of the FLOT1/FLOT2 heterodimeric complex. Related to Figure 4. **A)** Visualization of depth and diversity of the multiple sequence alignment (MSA). **B)** Predicted local distance difference test (plDDT) assessing local structural confidence of the different models. LDDT is a metric that evaluates local distance differences between all heavy atoms in a model, including validation of stereochemical plausibility. It ranges from 0 to 100, where 100 is the most confident. **C)** Five highest scoring ColabFold models for the FLOT1-FLOT2 heterodimer. Color coding of individual structural features is based on their plDDTs per residue. **D)** Inter-chain predicted aligned error (PAE), which aims to evaluate the position error at residue x, if the predicted and the true structures were aligned on residue y. It indicates the pairwise confidence of the respective prediction and ranks the models. **Abbreviations:** MSA: multiple sequence alignment; plDDT: predicted local distance difference test; PAE: predicted aligned error

**Figure S5:**
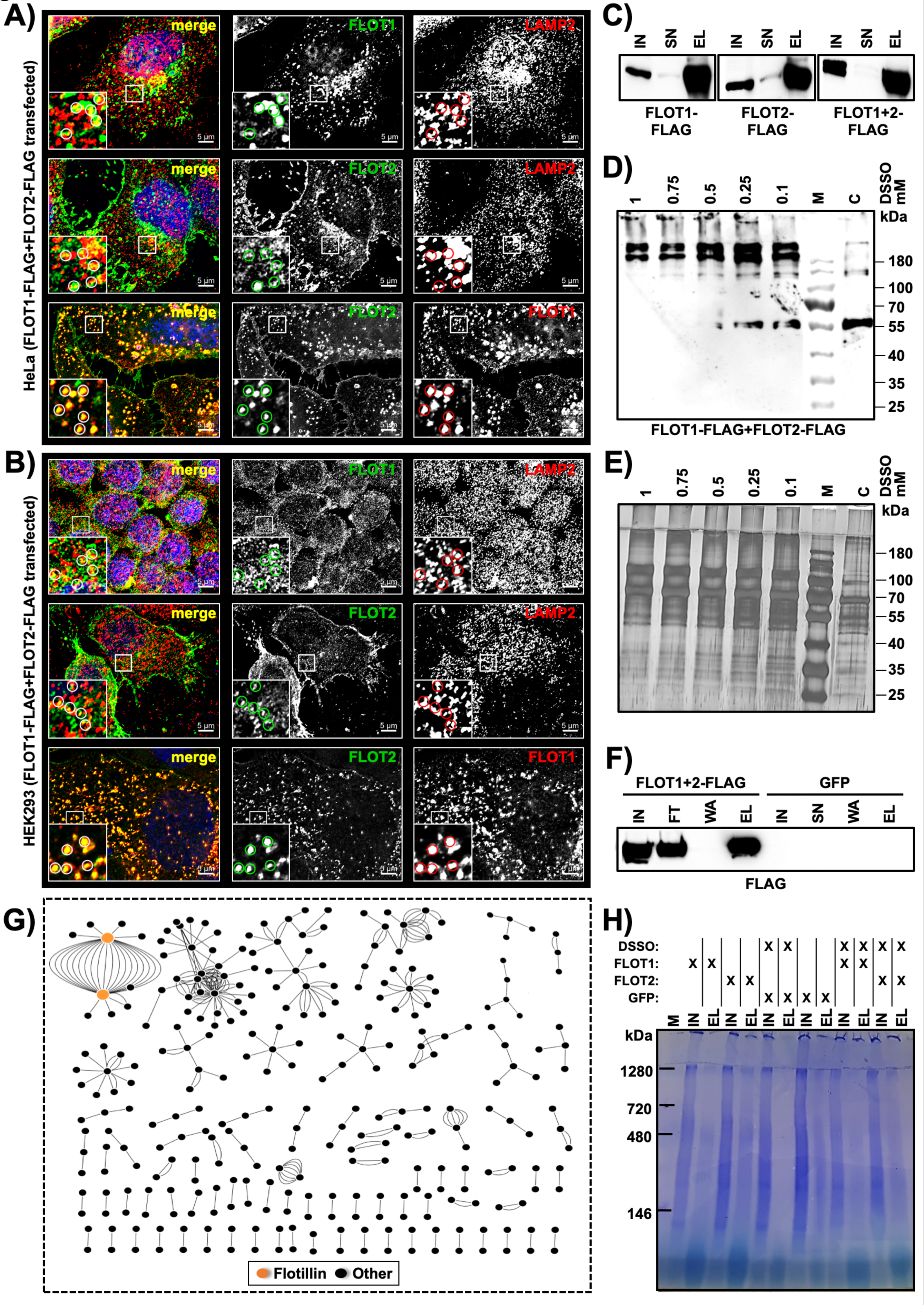
Investigation of higher order FLOT1/FLOT2 assemblies and XL-LC-MS/MS analysis of early endosomes. Related to Figure 5. **A/B)** Immunostaining analysis of HeLa and HEK293 cells transfected with FLOT1-FLAG+FLOT2-FLAG. Analyses were performed with anti-FLOT1 and anti-FLOT2 antibodies, co-localization with lysosomes was assessed through the lysosomal marker protein LAMP2. **C)** Western blot analysis for verification of FLOT1-FLAG, FLOT2-FLAG, and FLOT1-FLAG+FLOT2-FLAG expression and enrichment by FLAG-IP. **D)** Western blot analysis for determination of the optimal DSSO concentration for the cross-linking of SPIONs-enriched early endosomes from FLOT1-FLAG+FLOT2-FLAG transfected HEK293 cells. Early endosome-enriched fractions were cross-linked with the indicated amounts of DSSO at a protein concentration of 1 µg/µL followed by SDS-PAGE and anti-FLAG western blot. For each sample, 5 µg of protein were loaded to the gel. The non-cross-linked control sample (C) was treated with DMSO. Efficient cross-linking of FLOT1/FLOT2 complexes was judged by detection of the anti-FLAG band pattern. These analyses revealed that application of > 0.25 mM DSSO resulted in a near-quantitative loss of the FLOT1/FLOT2 monomers (∼55 kDa) with almost exclusive detection of higher molecular weight assemblies (>180 kDa) and increasing amounts of protein aggregates which did not enter the separation gel. **E)** Silver staining-based determination of the optimal DSSO concentration for the cross-linking of SPIONs-enriched early endosomes from FLOT1-FLAG+FLOT2-FLAG transfected HEK293 cells. Samples were generated and cross-linked as described in (D) followed by SDS-PAGE and silver staining of the gel. For each sample, 2.5 µg were loaded. The optimal DSSO concentration was determined to be 0.25 mM (final concentration), as the application of higher DSSO amounts resulted in reduced intensities in the low molecular weight region, indicating possible over cross-linking which could result in protein aggregates. Therefore, in accordance to the western blot results (see D), 0.25 mM DSSO was chosen. **F)** Verification of presence of FLOT1 and FLOT2 in SPIONs-enriched early endosomes from FLOT1-FLAG+FLOT2-FLAG overexpressing HEK293 cells prior to the XL-LC-MS/MS experiment. Loading amount: 10 % of each fraction. **G)** Interaction networks based on cross-links identified in the XL-LC-MS/MS dataset for DSSO-treated early endosome-enriched fractions from FLOT1-FLAG+FLOT2-FLAG-overexpressing HEK293 cells. In total, 324 inter-links were identified. FLOT1 and FLOT2 are highlighted in orange. The network was generated using Cytoscape. **H)** Coomassie staining of a BN-PAGE analysis of anti-FLAG-IP eluate fractions from HEK293 cells transfected with FLOT1-FLAG or FLOT2-FLAG. Input and eluate fractions were analyzed in an untreated and a DSSO-cross-linked state. Control cells were transfected with GFP. Loading amounts: IN: 50 µg, EL: 25 % of eluate fraction. **Abbreviations:** DSSO: disuccinimidyl sulfoxide; DMSO: dimethyl sulfoxide; PNS: post nuclear supernatant; M: marker; IN: input; SN: supernatant; FT: flow through; WA: wash; EL: eluate; C: control (no DSSO); IP: immunoprecipitation; BN-PAGE: blue native polyacrylamide gel electrophoresis; GFP: green fluorescent protein.

**Figure S6:**
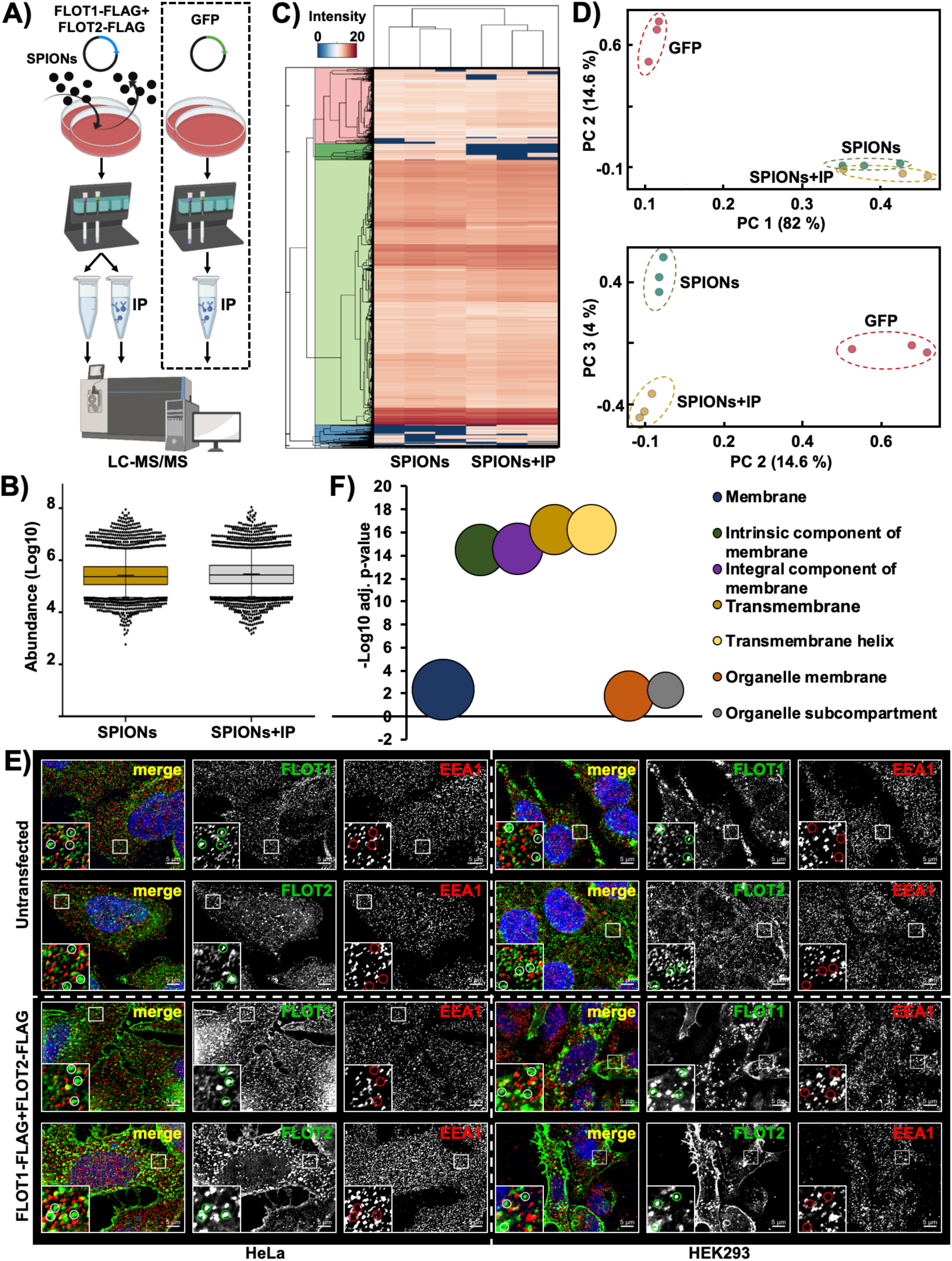
Determination of flotillin vesicular cargo. Related to Figure 6. **A)** Workflow for the enrichment of early endosomes by SPIONs and the subsequent enrichment of FLOT1-/FLOT2-positive early endosomal subpopulations by FLAG-IP. HEK293 cells were double-transfected with FLOT1-FLAG+FLOT2-FLAG, control cells with GFP. All samples were analyzed by LC-MS/MS using DIA. **B)** Analysis of protein abundances in DIA datasets of SPIONs-enriched early endosomes and FLAG-IP-enriched FLOT1-/FLOT2-positive early endosomes. Shown are combined values from three independent biological replicates, the median is indicated by a line, the average is marked with a “+”. **C)** Unsupervised clustering of protein abundances from three independent biological replicates for SPIONs-enriched early endosomes and FLAG-IP-enriched FLOT1-/FLOT2-positive early endosomes. Color coding correlates with the intensity of individual proteins. **D)** PCA for all SPIONs-enriched early endosomes, FLAG-IP-enriched FLOT1-/FLOT2-positive early endosomes, and GFP-transfected control cells. Four individual principal components were used (PC1 and PC2, as well as PC3 and PC4). **E)** Immunostaining analysis of HeLa and HEK293 cells. FLOT1, FLOT2, and EEA1 was analyzed both in untransfected cells as well as such transfected with FLOT1-FLAG+FLOT2-FLAG. A potential co-localization of FLOT1 and FLOT2 with the early endosome marker EEA1 was investigated. **F)** GO enrichment analysis applying STRING for proteins which are overrepresented in FLOT1-/FLOT2-positive early endosomes. Results from the GO-category cellular component, as well as the UniProt keywords transmembrane and transmembrane helix are shown. Bubble size correlates with the number of proteins assigned to an individual category. Shown are p-values corrected for multiple testing within each category using the Benjamini–Hochberg procedure (cut off < 0.05) representing the significance of enrichment (n=3). **Abbreviations:** SPIONs: superparamagnetic iron oxide nanoparticles; DIA: data independent acquisition; IP: immunoprecipitation; PCA: principal component analysis; adj.: adjusted; GO: gene ontology.

**Figure S7:**
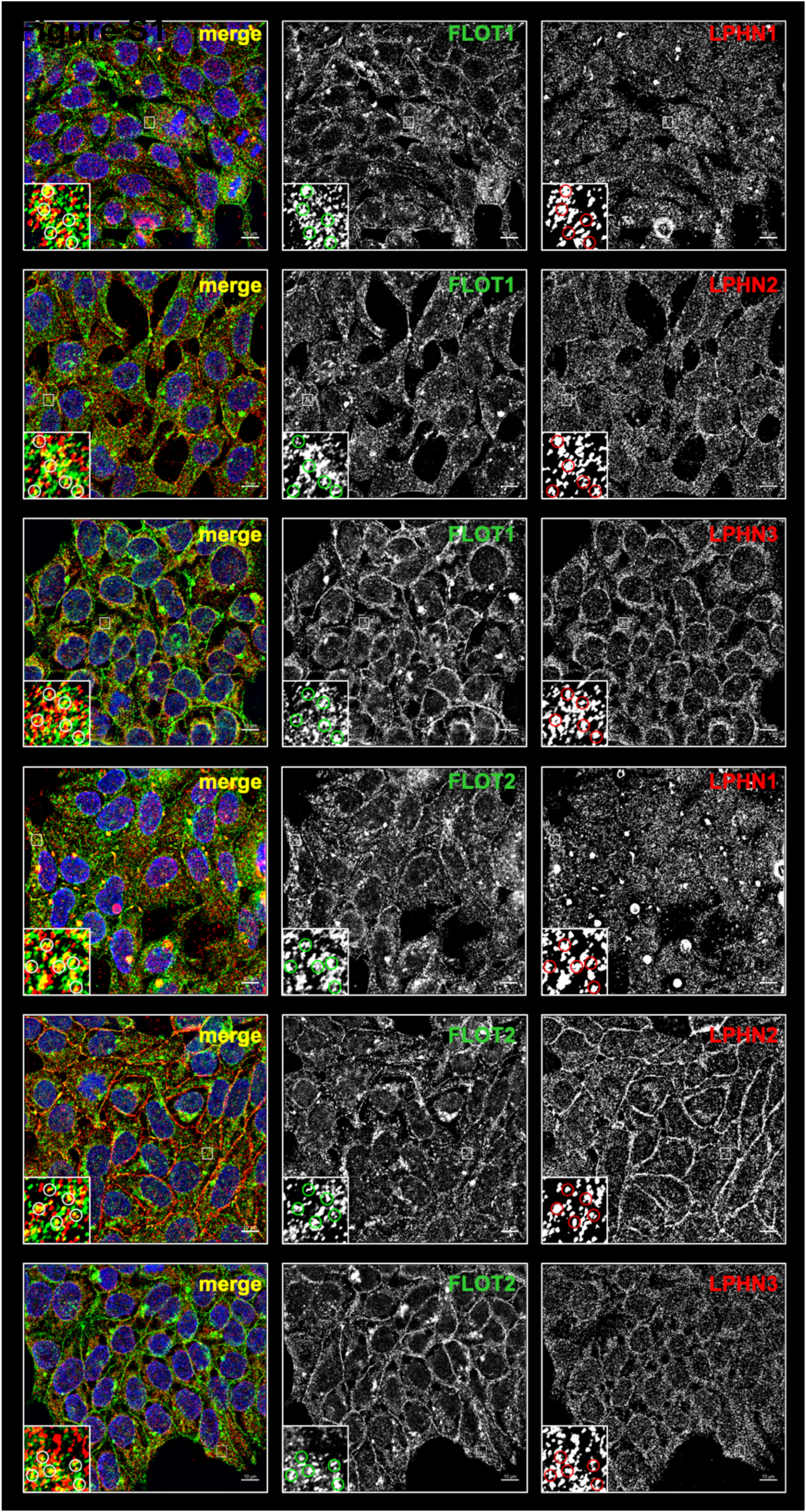
Latrophillins co-localize with FLOT1 and FLOT2 in punctate stainings. Related to Figure 7. Immunostaining analysis of HEK293 cells for FLOT1 and FLOT2 in combination with LPHN1, LPHN2, or LPHN3. Dependent on the combination of antibodies, different degrees of co-localization can be observed.

